# (A)symmetry in the Allele-Specific Chromosomal Structural Dynamics during Embryogenesis

**DOI:** 10.1101/2025.03.28.645915

**Authors:** Cibo Feng, Xiakun Chu

## Abstract

Embryonic stem cells (ESCs) lie at the heart of regenerative medicine and hold significant potential for treating various diseases. Understanding how the transcriptional landscape of ESCs is established during embryogenesis is therefore pivotal for deciphering the origins of life and advancing therapeutic strategies. Given the intrinsic connection between genome structure and function, chromosomal structural organizations during embryogenesis play a vital role in shaping gene expression patterns in ESCs. In this study, we employed a data-driven model and non-equilibrium molecular dynamics simulations to quantify large-scale chromosomal structural dynamics during embryogenesis. We focused on allelic differences and their impact on the interplay between chromosomal structure, dynamics, and function. Our results reveal that higher-order chromosomal structure, such as compartments and topologically associated domains (TADs), follow allele-symmetric developmental pathways, whereas the overall geometrical structures of chromosomes exhibit allele asymmetry, with the paternal and maternal chromosome undergoing monotonic and non-monotonic compactions, respectively. Despite these differences, the spatial distribution of chromosomal loci, particularly those rich in genes, adapts in an allele-symmetric manner. We propose that these chromosomal structural organizations during embryogenesis are intricately linked with epigenetic modifications and likely contribute to the transition from totipotency to pluripotency. Moreover, our findings suggest that allele asymmetry in chromosome structural dynamics during embryogenesis arises from long-range interactions, while short-range structures, particularly TADs, promote allele symmetry, a process associated with zygotic genome activation (ZGA). Overall, our findings provide theoretical insights into the dynamic establishment of the ESC genome during embryogenesis from the chromosomal structure perspective, and potentially lay the groundwork for further applications in the field.

## 1 Introduction

Embryogenesis marks the inception of life, initiated by the fusion of parental genomes to form a totipotent zygote ^[1]^. This fundamental event sets off a cascade of highly co-ordinated processes, including cell division, epigenetic remodeling, and the de novo establishment of chromatin architecture, collectively shaping the early developmental program ^[2–10]^. Underpinning these events is the three-dimensional (3D) organization of the genome, which not only provides a structural framework but also orchestrates the intricate regulation of gene expression. Consequently, deciphering how 3D chromosomal architecture evolves during early development is pivotal for advancing our understanding of cell fate decisions and the fundamental principles governing biological organization.

The 3D organization of the genome is non-random; rather, chromatin is intricately arranged into ordered architectures that are now routinely characterized by advanced techniques such as Hi-C ^[11, 12]^. This method, which captures genome-wide contact frequencies, provides a quantitative description of the average spatial chromosome organization at various scales ^[13]^. At the megabase level, chromatin is partitioned into topologically associated domains (TADs) that facilitate enhancer-promoter interactions critical for gene regulation ^[14–16]^. On a larger scale, genomic regions segregate into distinct compartments, reflecting transcriptionally active (euchromatin) and inactive (heterochromatin) domains ^[12, 17]^. Beyond these structural motifs, individual chromosomes occupy largely independent territories within the nucleus, where genes tend to localize at the interfaces between these domains to optimize accessibility to transcription factors ^[18]^. Together, these multiscale organizational features underscore the central role of 3D chromosomal architecture in regulating gene expression and cellular function.

Recent advances in low-input and single-cell Hi-C techniques have provided unprecedented insights into the reorganization of chromosomal structures during various stages of embryogenesis ^[19, 20]^. Evidence suggests that the paternal genome retains certain structural features inherited from sperm, whereas the maternal genome undergoes distinct remodeling dynamics throughout early development ^[19, 21]^. From both genetic and epigenetic perspectives, these findings indicate that the two allelic chromosomes exhibit asymmetric developmental behaviors. In contrast, the gradual establishment of chromatin accessibility appears to occur in a nearly symmetric manner ^[22, 23]^. Specifically, at the one-cell stage, the condensed sperm genome, which is initially packaged with protamines, is reassembled with histones ^[24]^. In the zygote, both parental genomes become significantly decondensed compared to their isolated gamete states, a crucial step for subsequent reprogramming ^[9]^. As development progresses, maternal proteins and mRNAs are gradually degraded, while the zygotic genome undergoes transcriptional activation ^[5, 7, 8, 25–28]^. Zygotic genome activation (ZGA) occurs abruptly, around the two-cell stage in mice, with two transcriptional surges that activate thousands of genes ^[29–35]^. Following ZGA, both allelic genomes undergo global DNA demethylation and gradually adapt their epigenetic modifications, structures, and functions during each cell cycle, including the progressive formation of TADs and compartments ^[5, 19]^. Over time, differences between the two allelic chromosomes become increasingly subtle. After several rounds of cell division, the first lineage specification event occurs, giving rise to the trophectoderm and inner cell mass (ICM) of the blastocyst. This transition ultimately leads to the formation of embryonic stem cells (ESCs) and the conversion of totipotency into pluripotency. While the timing of these key events has been broadly characterized, the precise relationship among 3D chromosomal architecture, epigenetic modifications, ZGA, and the totipotency-to-pluripotency transition remains largely unresolved ^[2–10]^.

One of the key challenges in studying chromosomal dynamics during embryogenesis lies in the complexity of the process, which involves multiple levels of regulation, including changes in DNA methylation, histone modifications, and protein factor binding ^[2–10]^. Additionally, the dynamic nature of chromosome organization makes it difficult to capture intermediate states during the transition from the zygote to ESC. Current experimental techniques are inherently limited by spatial and temporal resolution, preventing the acquisition of a complete developmental landscape. Moreover, conventional Hi-C experiments primarily provide ensemble-averaged chromosome structural information, which limits our understanding of molecular mechanisms and single-cell behaviors, particularly their impact on gene expression and embryonic development ^[36–41]^. Recent advances in computational modeling and molecular dynamics simulations have provided valuable tools for exploring chromosome structure and organization, enabling the study of chromosomal behavior at a molecular level and allowing for high-temporal-resolution predictions of structural changes ^[42–49]^. In order to describe the chromosome structure dynamics accurately, a variety of strategies, classified into data-driven and physics-based models, are applied to optimize the models, acquiring the relationship between structures, dynamics and functions of chromosomes ^[50]^.

Here, we employ a landscape-switching model combined with coarse-grained molecular dynamics simulations ^[42–49]^ to investigate chromosome structural dynamics during embryogenesis. In our approach, the landscape-switching model is applied independently to the two allelic chromosomes, allowing us to monitor their respective developmental behaviors throughout early embryogenesis. First, we demonstrate that our model accurately reproduces the experimentally measured intermediate chromosomal states. Subsequently, the continuous trajectories generated reveal a comprehensive dynamic process from the perspective of chromosomal structure. Notably, higher-order chromosome structures, such as compartments and TADs, are established in a nearly symmetric fashion, whereas the overall geometric shape of the chromosomes evolves asymmetrically, with the paternal chromosome exhibiting a monotonic change and the maternal chromosome displaying a non-monotonic behavior. Despite these geometric differences, the relative positional distribution of chromosomal loci, particularly genes, remains symmetric. Furthermore, our analysis indicates that the maternal chromosomal non-monotonic geometric evolution arises predominantly from long-range structural reconfigurations, while allelic symmetry is maintained within short-range structures, including intra-TAD and short-range inter-TAD regions. By characterizing these chromosome structural features, we establish a connection between TAD organization and ZGA, and elucidate the relationships among compartmental organization, epigenetic modifications, and the totipotency-to-pluripotency transition. Overall, our study provides molecular-level insights into the dynamic reorganization of chromosome structures during embryogenesis, reveals distinct developmental behaviors across different structural scales, and uncovers the underlying mechanisms linking chromosomal structure, dynamics, and function. These findings offer a theoretical framework that can be validated in future experiments and may inform potential applications in congenital disease treatment and stem cell therapy.

## 2 Results and Discussion

### 2.1 Quantified Dynamical Chromosome Structural Organization during Embryogenesis

To investigate the large-scale reorganization of chromosomal structures during embryogenesis, we applied a landscape-switching model combined with coarsegrained molecular dynamics simulations ^[42–49]^. Our approach consists of two main steps. First, we generated 3D chromosomal structure ensembles for the initial (zygote) and final (ESC) cell states using energy landscapes derived from the maximum entropy principle ^[51–53]^ and trained with corresponding Hi-C data ^[19]^. Given the early allelic differences, separate ensembles were constructed for the paternal (ZP) and maternal (ZM) zygotes, whereas in ESC the chromosomes are treated as structurally indistinguishable (Figure 1A, Figure S1-S3). Second, the developmental process for each allele was modeled independently. To simulate the cell-state transition, the chromosome structures in the ZP (or ZM) energy landscape (Figure S4) were excited and then allowed to relax toward the ESC state via an energy landscape switch, producing a series of transition trajectories. The chromosome structures sampled at each time point form a quasi-ensemble that captures the temporal evolution of chromosomal organization. Subsequently, Hi-C-like contact maps (see supplementary information for details) were calculated from these time-course ensembles (Figure 1A), revealing a smooth, continuous reorganization process from the initial to the final state.

**Figure 1:**
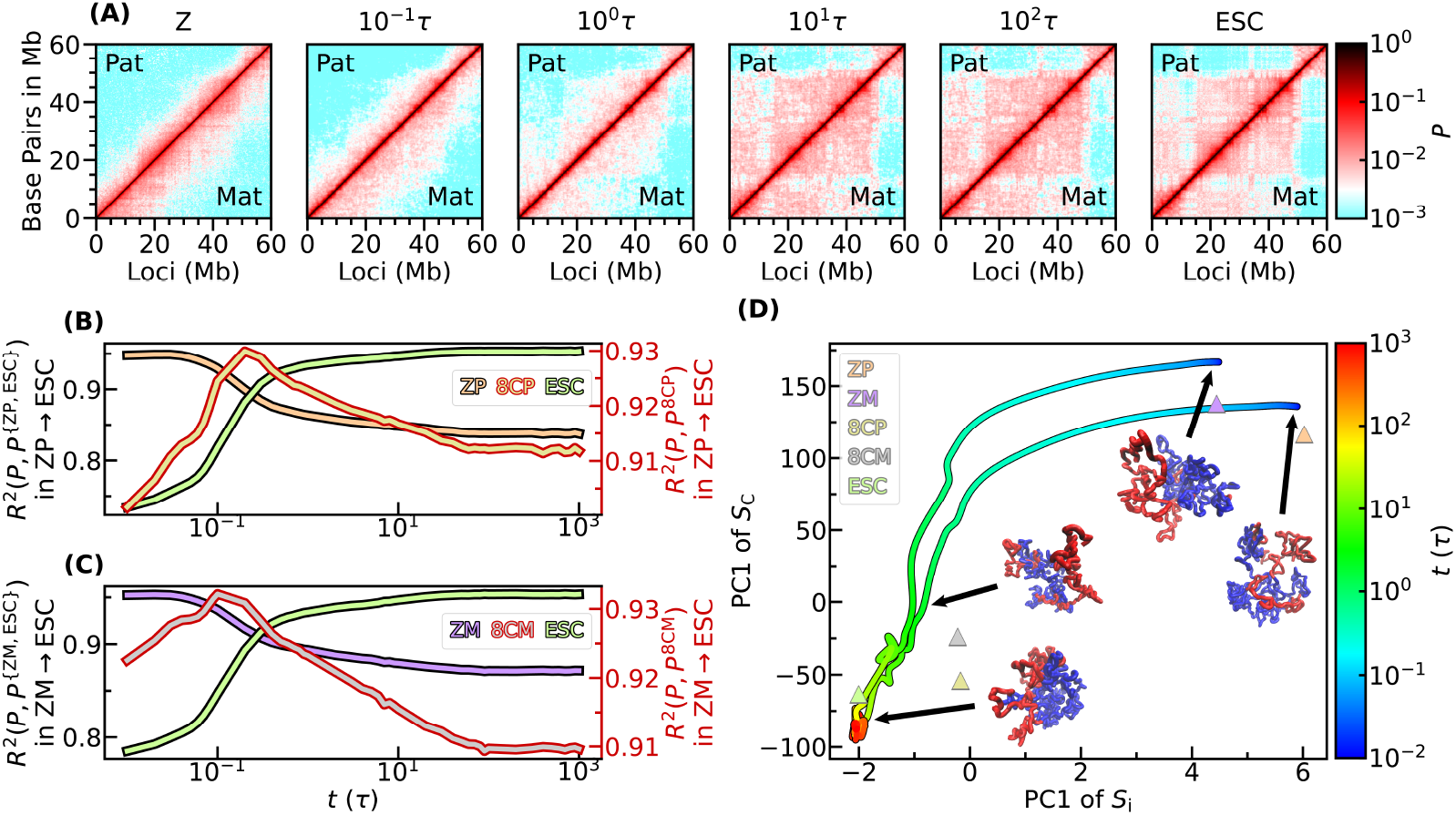
Simulated chromosome states transition processes and comparison with experimental intermediate states. (A) Time-course chromosome contact maps during embryogenesis. The upper left triangles are those of paternal chromosome and lower right triangles correspond to maternal one. The first and last figures, as initial and final states, are from Hi-C experiments of zygote and embryonic stem cell (ESC). The middle ones, as intermediate states at 0.1*τ*, 1*τ*, 10*τ*, and 100*τ*, respectively, are predicted by landscape switching model (*τ* is the reduced time unit). (B) Structural similarity between chromosomes in transition and reference cells: paternal zygote (ZP), paternal 8-cell stage (8CP), and ESC. The curve referring to 8CP is aligned to the right y-axis. (C) Similar to (B), but for maternal allele. The reference cells are maternal zygote (ZM), maternal 8-cell stage (8CM), and ESC. (D) Development pathways during embryogenesis of both allelic chromosomes projected to the first principal component (PC1) of insulation score *S*_i_ and enhanced contact map *S*_C_. The triangles represent the values of reference cells.

To validate the simulated trajectories, we compared the Hi-C contact maps *P* generated from simulations with those obtained from experiments ^[19, 20]^. Specifically, we used ensemble Hi-C data from the 8-cell stage (8CP and 8CM for the paternal and maternal allelic chromosomes, respectively) as a reference for comparison. To quantify the similarity between simulated and experimental chromosome structures, we calculated the coefficient of determination, *R*^2^(*P, P*^ref^ ) (see supplementary information for details), where a higher *R*^2^ value indicates greater similarity, with a maximum value of 1 representing an exact match (Figure 1B-C). As expected, during the transition, the similarity between the chromosome structure and its initial state (zygote) gradually decreases, while its similarity to the final state (ESC) increases. Notably, the similarity to the 8-cell stage first increases and then declines, suggesting that the simulated transition captures an intermediate chromosomal state resembling the 8-cell stage. This trend is further supported by analyses of chromosomal compartments and TAD structures, quantified using the enhanced contact probability matrix *S*_C_ and the insulation score *S*_i_ (Figure 1D, see the supplementary information for details). Principal component analysis (PCA) of these structural features reveals that the developmental trajectories of both allelic chromosomes pass through the 8-cell stage. Meanwhile, the evolution of two allelic chromosomes merges near here, indicating the progressive loss of allelic specificity during embryogenesis. Additionally, our results suggest that chromosomal reorganization initially occurs predominantly at the TAD level, before shifting toward compartment-level restructuring. This hierarchical adaptation, where local chromosome structures (e.g., TADs) stabilize before long-range (e.g., compartment) organization, aligns with general principles of polymer dynamics ^[54, 55]^ and has been widely observed in studies of 4D chromosome dynamics ^[49]^.

### 2.2 Gradual Formation of High-order Chromosomal Structures by Scale-dependent Dynamics during Embryogenesis

In early embryogenesis, the initially loose organization of the zygote undergoes dramatic compaction and reorganization ^[4, 7, 8, 10, 19, 20]^. At this stage, distinct higher-order chromosomal structures, such as compartments ^[19, 21, 56]^ and TADs ^[19, 21, 57]^, are not yet established. As development proceeds, these structures emerge progressively. To quantitatively elucidate the formation mechanisms of compartments and TADs, we calculated the compartmentalization strength *A*_C_ and insulation strength *A*_i_. Specifically, *A*_C_ is derived from the mean interaction strengths both within (AA and BB) and between (AB) compartments, as captured by the enhanced contact probability matrix 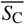^[12,58,59]^, whereas *A*_i_ is defined as the depth of the mean boundary insulation score 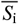^[19,58,60]^ (Figure 2, see supplementary information for details).

**Figure 2:**
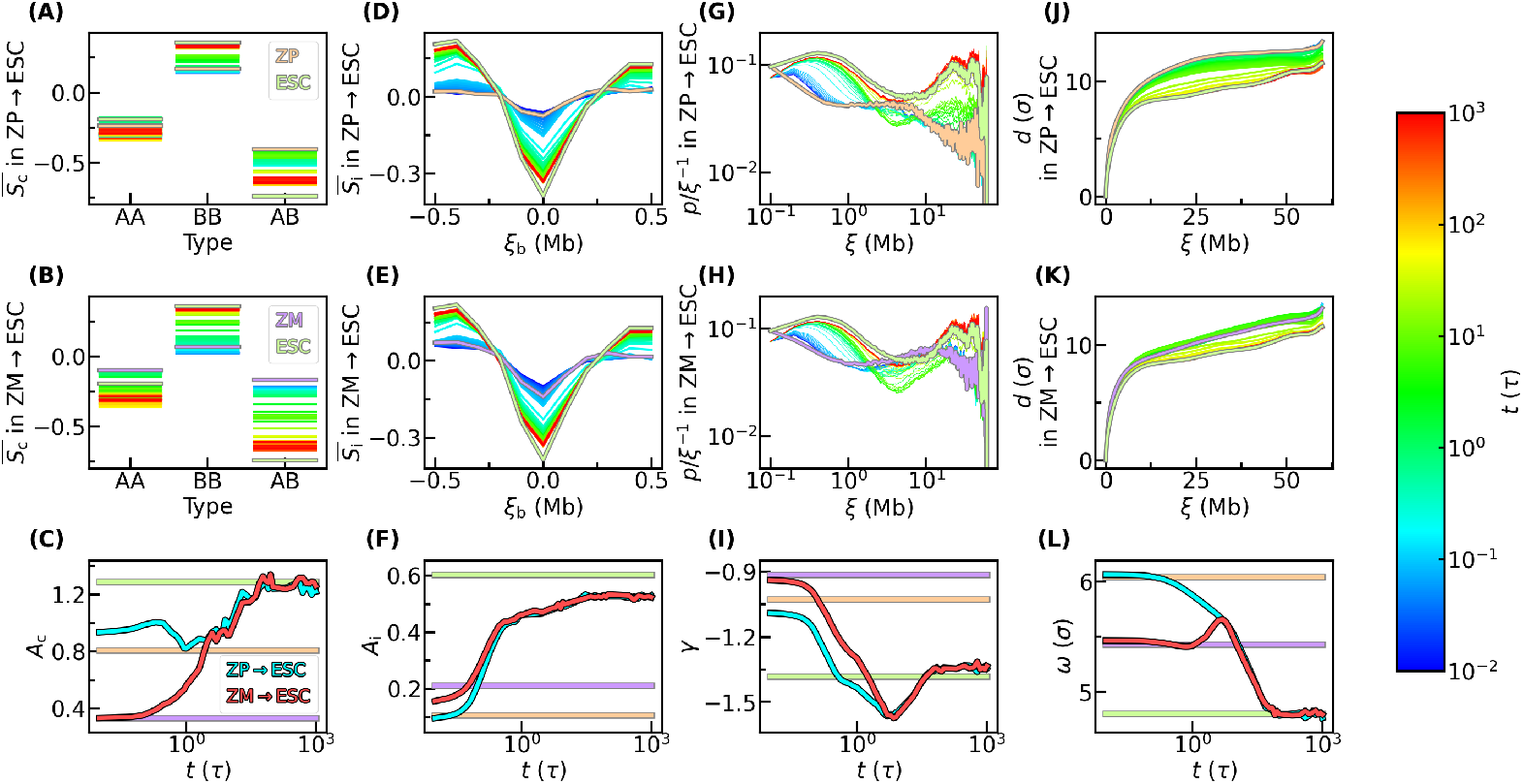
Evolution of higher-order chromosome structures during embryogenesis. (A) Evolution of the mean interaction strength between (AB) and within (AA and BB) compartments of paternal chromosome. The curves with gray border represent the values of reference cells. (B) Similar to (A), but for maternal one. (C) Compartmentalization strength change over time. The lines with gray border represent the values of reference cells. (D) Evolution of mean boundary insulation score of paternal chromosome. The curves with gray border represent the curves of reference cells. (E) Similar to (D), but for maternal one. (F) Insulation strength change over time. The lines with gray border represent the values of reference cells. (G) Evolution of the *p* decay curve of paternal chromosome, normalized by a standardized function *p*_0_ = *l*^−1^. The curves with gray border represent the values of reference cells. (H) Similar to (G), but for maternal one. (I) Power exponents characterizing polymer state change over time. The lines with gray border represent the values of reference cells. (J) Evolution of the *d* v.s. *ξ* curve of paternal chromosome. The curves with gray border represent the values of reference cells. (K) Similar to (J), but for maternal one. (L) Parameters of *d*_*ξ*_ v.s. *ξ* curves change over time. The lines with gray border represent the values of reference cells.

For our focused chromosome segment, the data indicate that interactions are predominantly enriched within compartment B, whereas those within compartment A are comparatively weaker, and the interactions between compartments are the least robust. As development proceeds, contacts within compartment B become further strengthened, while those within compartment A and across compartments gradually weaken (Figure 2A-B). Consequently, the overall compartmentalization is enhanced over time (Figure 2C). Owing to partial inheritance from sperm ^[7, 19, 21, 56]^, the paternal chromosome begins with a higher degree of compartmentalization; however, its consolidation occurs more gradually. Around 10*τ*, where *τ* is the reduced time unit in simulations, the compartmentalization strength *A*_C_ of the maternal chromosome catches up to that of the paternal chromosome (Figure 2C), after which both allelic chromosomes develop in synchrony. Notably, the progression of compartmentalization in the paternal chromosome is characterized by subtle fluctuations (Figure 2C), consistent with experimental observations ^[19]^. Overall, the developmental trajectories of the two allelic chromosomes exhibit remarkably similar, allele-symmetric trends.

Turning our attention to TAD organization, we observed that inter-TAD interactions decrease gradually while intra-TAD contacts become increasingly enriched (Figure 2D-E). This trend indicates a progressive consolidation of TAD boundary insulation during embryogenesis. Notably, although slight allele-specific differences exist, both ZP and ZM exhibit similarly low insulation strengths *A*_i_, with their TAD dynamics converging as early as 0.3*τ*. In contrast to compartment formation, TAD consolidation proceeds at a faster rate (Figure 2F), in agreement with experimental observations ^[7, 19, 57]^. This uncoupling between the dynamics of compartment segregation and TAD establishment underscores that distinct mechanisms underlie their formation ^[9, 57, 61]^, and even a certain antagonizing relationship between them ^[6, 62, 63]^.

Overall, the TAD developmental trajectories are remarkably allele-symmetric, highlighting a unified local organizational strategy during early development.

To capture the scale-dependent dynamics of chromosomal compaction and reorganization, we calculated the mean contact probability *p* between chromosomal loci separated by a linear distance of *ξ* Mb (see the supplementary information for details). These curves are normalized by a reference function *p*_0_ = *ξ*^−1^, enabling clearer comparisons across them. As expected, for distances from 0 to 2 Mb (the typical scale of TADs), *p* changes monotonically, consistent with the gradual strengthening of TADs. In contrast, beyond 2 Mb, despite the overall increase in compartmentalization, the *p* curves display a complex pattern with both monotonic and non-monotonic segments, suggesting that chromosomal compaction and compartmentalization are uncoupled processes (Figure 2G-H). To further characterize the polymeric state of the chromosome, we fitted the relationship between *p* and *ξ* (in the range of 0.5–7 Mb) to a power law of the form *p* ∝ *ξ*^*γ* [44,64,65]^ (see supplementary information for details). Here, a higher *γ* indicates a more compact polymer conformation. As illustrated in Figure 2I, the evolution of *γ* follows an allele-symmetric but non-monotonic trajectory. This can be speculated as the competition between compartment and TAD ^[66]^, as they work at different stages and distance scales. Initially, the decrease in *γ* coincides with the rapid establishment of TADs, likely because enhanced local contacts (0.5–1 Mb) diminish the relative contribution of longer-range interactions (1–7 Mb). Later, as TAD formation plateaus, the subsequent increase in *γ* is primarily driven by the strengthening of contacts within compartment B, which spans longer genomic segments than compartment A, reflecting a phase dominated by compartmentalization. This dynamic interplay is reminiscent of the chromosome structural transitions observed during mitotic exit ^[42]^, where non-monotonic changes in chromosomal architecture can also be attributed to the competition between compartment formation and TAD consolidation.

Characterization of long-range chromosomal organization is facilitated by analyzing the fractal relationship between spatial and sequence distances. In our study, the paternal chromosome exhibits a monotonic shift, whereas the maternal chromosome displays a non-monotonic behavior (Figure 2J-K), highlighting an allelic asymmetry. Considering the power law of these curves in long distance region is around *d* ∼ *ξ*^0.2^ (Figure S5A-B), by fitting them to a power law *d* = *ωξ*^0.2^ (see supplementary information for details), we parameterize the curves with the coefficient *ω*, where a higher *ω* indicates a more relaxed chromosomal conformation. Notably, the maternal chromosome shows a significant expansion peak at around 10*τ*, in contrast to the steady behavior of the paternal chromosome (Figure 2L). This over-expansion in the maternal allele occurs during compartmentalization, although it does not appear to influence the overall progression of compartment formation.

To further elucidate the kinetics and dynamics of chromosomal reorganization across different genomic scales, we calculated the compactness *P*_𝕃_, folding progress *Q*_𝕃_, and their corresponding half-lives, with 𝕃 representing a specific genomic distance interval. Here, *Q*_𝕃_ measures the chromosome structural similarity to the final state (ESC), analogous to metrics used in protein folding. Our analysis reveals that the paternal chromosome exhibits non-monotonic compactness predominantly in the 2–6 Mb range, whereas the maternal chromosome shows a broader non-monotonic pattern that even though also gradually diminishes at larger scales (the upper right triangle of Figure S6). Additionally, compaction within the 0–2 Mb range proceeds more rapidly compared to longer distances; beyond 2 Mb, the compaction rates become largely scale-independent. In contrast, the paternal chromosome’s folding progress remains relatively smooth, while the maternal chromosome demonstrates increasing non-monotonicity with distance (the lower left triangle of Figure S6). This disparity suggests that chromosomal expansion is closely linked to folding dynamics. Furthermore, the scale dependency of the folding rate spans up to 6 Mb, slightly wider than that observed for compactness. A direct comparison of these two metrics indicates that, except in the 2–6 Mb region where folding precedes compaction, the processes occur synchronously (the diagonal of Figure S6). Overall, these findings underscore that chromosomal compaction and folding are intricately coupled during embryogenesis.

In further analyses, we investigated the heterogeneity of chromosomal reorganization at the chromosome locus level by calculating compactness *P*_*i*𝕃_ and folding progress *Q*_*i*𝕃_ metrics formed by each chromosome locus *i* across various genomic distance intervals L (see supplementary information for details). To facilitate kinetic comparisons, the transition processes were normalized (see supplementary information for details). Our results reveal marked heterogeneity in chromosomal behavior, particularly at intermediate distances (Figure S7A-H and Figure S8A-H). Notably, the heterogeneity in compactness is substantially greater than that observed in folding progress. Quantitatively, the mean half-lives *T*_1*/*2_ of these transitions increase with genomic distance, although this scale dependency of compactness diminishes for distances greater than 6 Mb, in accordance with the previous results. Moreover, the ratio of the second-order moments, 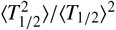 , which quantifies locus-level heterogeneity, peaks at intermediate length scales (Figure S7I and Figure S8I). The lower heterogeneity at both short and long distances aligns with patterns observed during mitotic exit ^[42]^. However, the high heterogeneity at intermediate distances highlights the complex interplay between TAD formation and compartment segregation in modulating chromosomal architecture during embryogenesis.

### 2.3 Allele-(A)symmetric Change of Geometry Shape and Spatial Adaptation of Chromosomal Loci

To comprehensively capture the evolution of chromosomal geometry during embryogenesis, we quantified detailed shape parameters, including the lengths of the principal inertial axes (*R*_PA1_, *R*_PA2_, and *R*_PA3_), the radius of gyration (*R*_g_), and aspherical metrics (*Δ* and *S*) (see supplementary information for details). Notably, the ZP chromosome is significantly larger than the ZM one, consistent with experimental observations of a more compact contact map for the ZM chromosome ^[19–21]^. Although substantial heterogeneity exists among individual chromosome ensembles (Figure 3A-B, D-E, and Figure S9), the ensemble-average trajectories are clearly discernible. Specifically, the overall size of the paternal chromosome decreases monotonically throughout development, both in total and along each principal axis, whereas the maternal chromosome exhibits a pronounced non-monotonic change (Figure 3 and Figure S9A-D). In the maternal case, an initial slight decrease, primarily in the shorter axes, is followed by a notable expansion in the longer axes, and finally a significant contraction. This allele-specific behavior suggests distinct regulatory mechanisms modulating chromosomal geometry during early development (Figure S9A-C).

**Figure 3:**
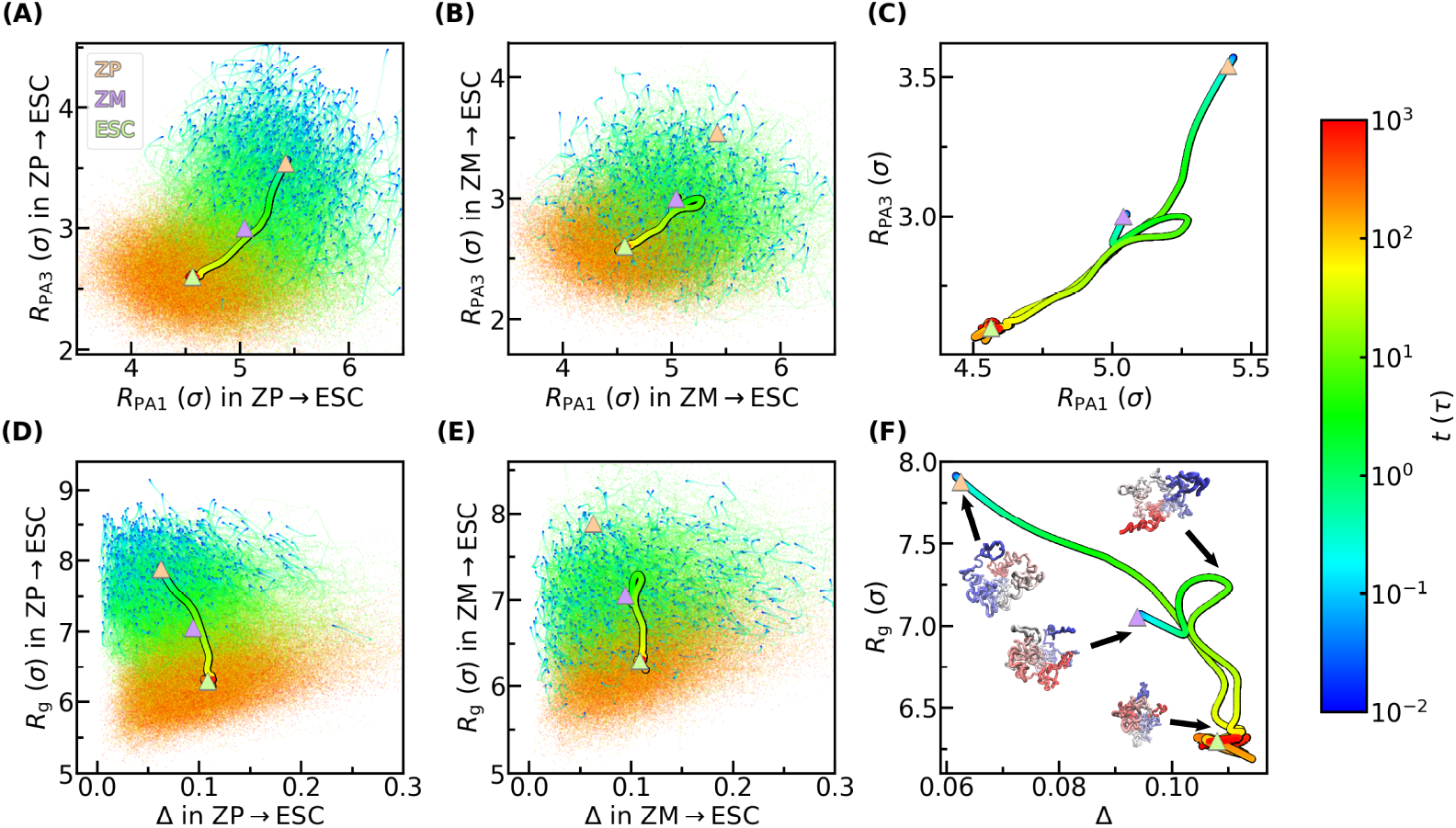
Change of chromosome geometry shape during embryogenesis. (A) Change of the longest and shortest inertial principal axes length (*R*_PA1_ and *R*_PA3_) of paternal chromosome. The fine curves represent the values of each single chromosome. The thick curve represents the values of ensemble average. The triangles represent the values of reference cells. (B) Similar to (A), but for maternal one. (C) The combination of (A) and (B), excluding curves of single chromosomes. (D-E) Similar to (A-C), but for aspherical degree (*Δ*) and radius of gyration (*R*_g_).

The expansion of the maternal chromosome may be linked to global DNA demethylation ^[5]^, a process also known to promote telomere elongation in early embryos ^[67, 68]^. Moreover, our simulations reveal that compartment A undergoes greater expansion than compartment B (Figure S10), which is consistent with experimental evidence indicating that DNA demethylation predominantly affects compartment A ^[21]^. This further substantiates the conjecture. In contrast, the subsequent compaction observed in both allelic chromosomes appears to be associated with de novo DNA methylation ^[69, 70]^. However, it remains unclear why the paternal chromosome does not exhibit a similar expansion. One plausible explanation is that allele-specific histone modifications occurring after ZGA, particularly changes in H3K27me3, H3K9me3, and H3K64me3, may play a role. For instance, H3K27me3 is lost from the maternal chromosome post-ZGA while it is maintained in the paternal chromosome ^[3, 71]^. Given that the absence of H3K27me3 is associated with gene activation and a more relaxed chromatin state, this could contribute to the maternal expansion. Likewise, a decreasing proportion of maternal-specific H3K9me3 relative to the paternal form ^[72]^, along with the global erasure of H3K64me3 ^[73]^, supports the notion of a more relaxed organization in the maternal allele. An additional mechanism to consider is that a preferential transfer of histones from the maternal to the paternal chromosome needed for protamine replacement, could result in a relative histone deficiency in the maternal genome, further contributing to its expansion. Intriguingly, the pronounced peak in maternal chromosome size seems to mark a critical transition point, after which the sizes of the two chromosomes converge rapidly. This implies the non-monotonic behavior of the maternal chromosome may contribute to the allelic merging as fast as possible. Notably, the primary changes in chromosome size are nearly synchronous with alterations in compartmentalization, but lag behind TAD modifications (Figure S9A-D and Figure 2I), underscoring a tight coupling between chromosomal geometry and compartment organization.

Meanwhile, the overall reduction in chromosome size leads to an increased asphericity, particularly for the paternal chromosome (Figure S9E). This observation suggests that the chromosome adopts a more tube-like conformation, as indicated by the asphericity parameter *S* (Figure S9F), becoming progressively more slender and elongated throughout embryogenesis. Notably, when comparing the lengths of the first (longest) and third (shortest) principal axes, we find that the initially shortest axis contracts more dramatically during the early stage, driving the marked increase in asphericity. In the later stage, the contraction becomes more uniform across the axes, leading predominantly to overall size reduction rather than further shape distortion (Figure 3D-F and Figure S9E). In other words, the contraction of the shortest axis of the chromosome in the early stage mainly increases the asphecial degree, while the contraction of the two axes of the chromosome in the same proportion in the late stage mainly reduces the overall size.

The spatial organization of chromosomal loci, particularly genes, plays a pivotal role in regulating gene expression ^[74, 75]^. To elucidate the distribution of chromosomal loci, we calculated the radial distance *r* of each locus from the chromosomal center of mass and analyzed these distributions for loci located in compartment A, compartment B, and genes (Figure S11). Because these distributions are inherently influenced by overall chromosome size, we normalized them by the radius of gyration *r/R*_g_ of the chromosome (Figure 4). Intriguingly, despite the markedly different geometric evolution of the two allelic chromosomes, the normalized spatial distributions of chromosomal loci evolve in a highly allele-symmetric fashion. Specifically, as embryogenesis progresses, loci in compartment A increasingly localize toward the periphery of the chromosome, whereas loci in compartment B shift toward the center. Consistent with this, genes also tend to relocate closer to the surface.

**Figure 4:**
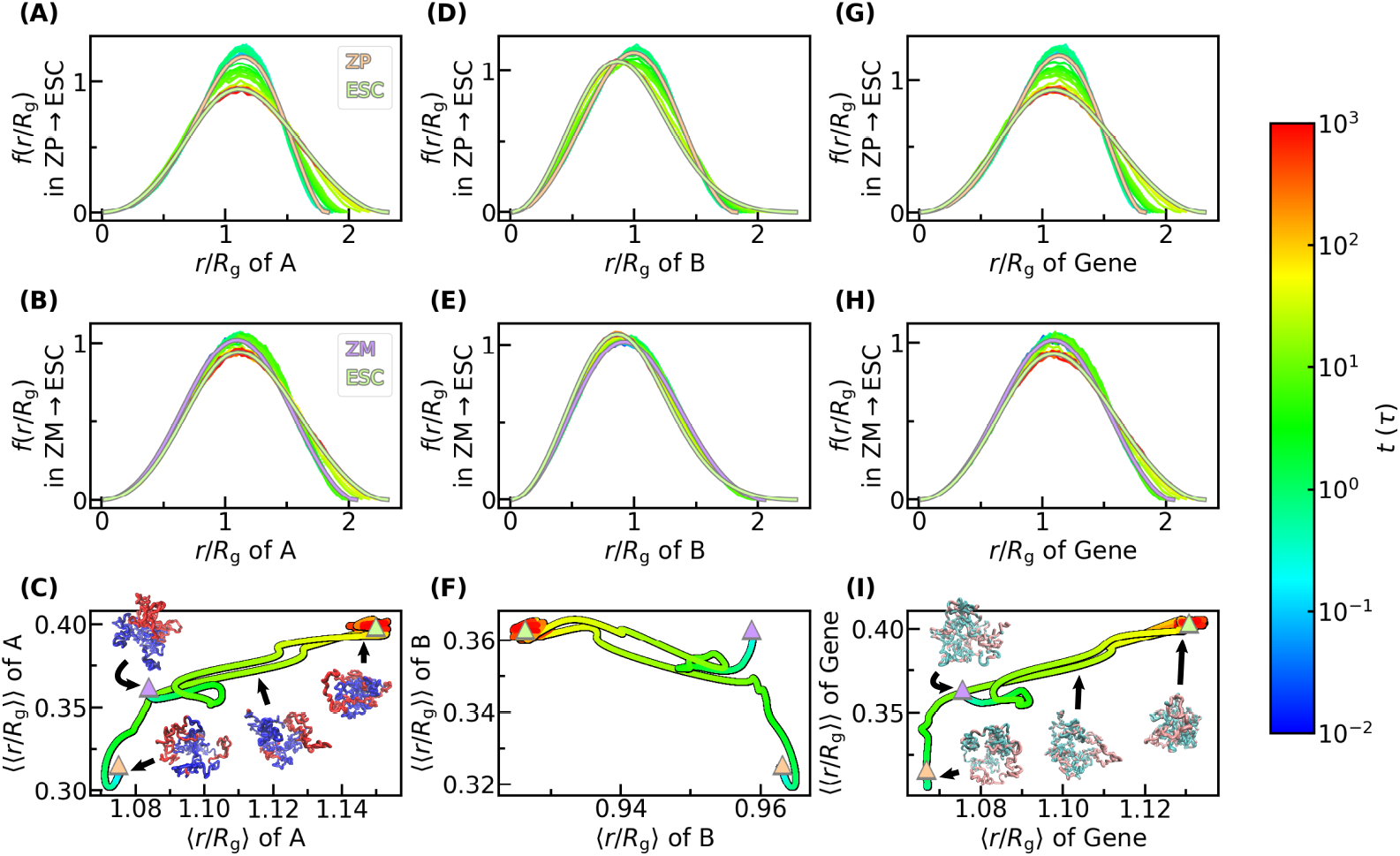
Adaptions of loci distribution in chromosomes during embryogenesis. (A) Adaptions of distribution of relative radial position of chromosomal loci in compartment A for the case of paternal chromosome. The curves with gray border are the distributions of reference cells. (B) Similar to (A), but for maternal one. (C) Mean and standard deviation of the relative radial position of chromosomal loci. The triangles with gray border represent the values of reference cells. (D-F) Similar to (A-C), but for loci in compartment B. (G-I) Similar to (A-C), but for genes.

The adjustment in chromosomal loci positioning occurs predominantly during the late stages of embryogenesis (10*τ*–100*τ*) (Figure S12E-F), coinciding with the convergence of the overall chromosomal geometries following initial allele-specific adaptations (Figure 3 and Figure S9). Notably, even the extensive establishment of compartmentalization in the maternal chromosome during 0.1*τ*-10*τ* does not significantly alter the normalized compartmental distributions (Figure 2I, Figure 4A-F, and Figure S12E-F). These observations imply that ZGA, which occurs early, may not critically depend on the global compartment structure or gene distribution. Indeed, previous experiments have indicated that reducing compartmentalization has minimal impact on ZGA ^[76]^. Thus, the consolidation of compartments and the repositioning of gene loci may primarily serve to inhibit the expression of a subset of genes in compartment B rather than directly activating most genes in compartment A, thereby facilitating the transition from totipotency to pluripotency ^[2, 77–82]^. This conclusion is further supported by the delayed onset of de novo methylation ^[2, 5, 69, 70, 83]^ and the maturation of repressive histone modifications ^[2, 5, 71, 72, 84, 85]^, relative to active epigenetic marks ^[2]^. Collectively, these data suggest that the functional relationship between 3D chromosomal architecture and ZGA is primarily mediated by TAD dynamics, whose temporal pattern coincides with the onset of ZGA ^[9]^. This relationship is also suggested in experiment ^[81, 86–91]^. For instance, experimental evidence indicates that the key ZGA gene *Zscan4* is expressed only when TAD structures are not fully established ^[87, 88]^.

Interestingly, even after normalizing for overall chromosome size, the trajectories of the chromosomal loci distribution profiles exhibit behavior that mirrors the evolution of the shape parameters. In particular, the maternal chromosome follows a complex, circular path before converging with the paternal chromosome (Figure 4C,F,I). This suggests the pathway relationship between the two allelic chromosomes is similar at different levels.

### 2.4 Varying TAD-based Behaviors in Allelic (A)Symmetrical Chromosome Structural Dynamics during Embryogenesis

To elucidate the relationship between TAD organization and functional regulation, we examined the interaction patterns associated with TADs. When considering the radius of gyration 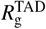 of TADs, the distributions evolve monotonically over time for both allelic chromosomes (Figure 5A-C), consistent with the behavior observed in the *p* v.s. *ξ* curve for distances between 0.1 and 2 Mb. In contrast, the gap distance between neighboring TADs 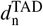 , defined as the distance between the mass centers of adjacent TADs after subtracting their radii, exhibits non-monotonic changes over time for both allelic chromosomes (Figure 5D-F), which corresponds to the 2–10 Mb range of the *p* v.s. *ξ* curve. For larger scales, we evaluated the gap distance between non-consecutive TADs (*d*^TAD^) by plotting the gap distance between TADs separated by *ξ*^TAD^ (Figure 5G-H). Unlike the typical power-law behavior of fractal curves, this relationship follows a logarithmic law (Figure S5E-F). Notably, for *ξ*^TAD^ greater than 10, approximately corresponding to regions beyond 10 Mb, the maternal chromosome shows a marked non-monotonic shift in the *d*^TAD^ curve, whereas the paternal one displays only minor non-monotonicity. This difference corresponds to the allele-asymmetry. When fitting these curves to *d*^TAD^ = *Ω*ln*ξ*_TAD_ (see supplementary information for details) and monitoring the change of parameter *Ω*, with a higher value corresponds to a more relaxed inter-TAD space, a minor non-monotonic peak in the parameter *Ω* appears at approximately 0.3*τ* for both allelic chromosomes, representing a symmetric component, similar to that in 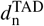. However, the maternal chromosome also exhibits a pronounced non-monotonic peak at around 5*τ*, reflecting allele-asymmetric behavior. These results suggest that allelic asymmetry predominantly arises from variations in long-range inter-TAD organization, while intra-TAD and short-range inter-TAD structures contribute in a symmetric fashion. Interestingly, the symmetric component of long-range inter-TAD distance expansion also occurs early, coinciding with the condensation of TADs and expansion of short-range inter-TAD distance. Although this simultaneous TAD condensation and inter-TAD expansion may appear contradictory, these processes likely act in concert to promote ZGA: the former enhances cis-contacts, such as enhancer-promoter interactions, whereas the latter insulates against improper contacts, facilitating the establishment of an ordered gene regulatory network and allowing trans-acting factors better access to the chromosomal interior.

**Figure 5:**
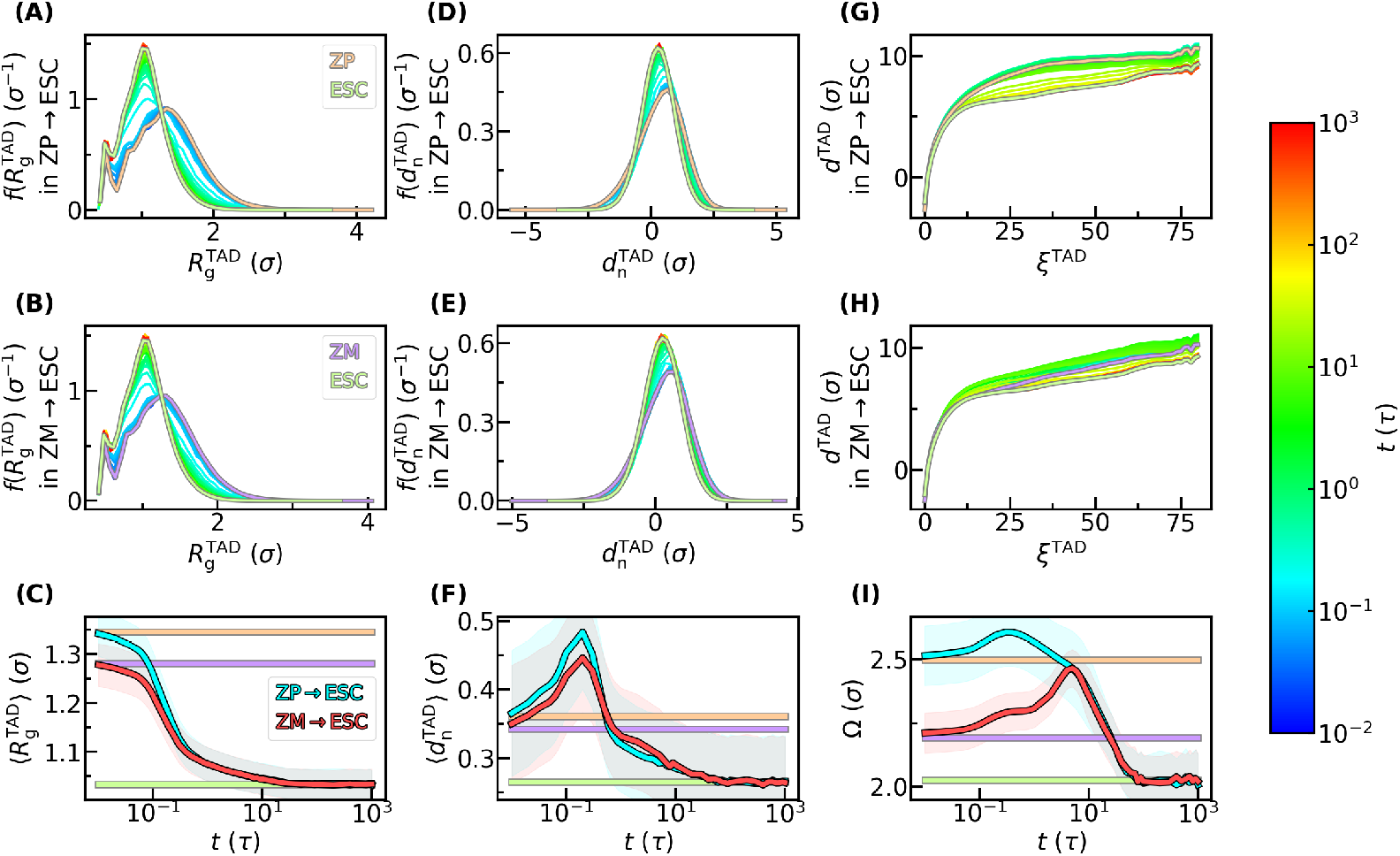
Structural characteristics within and between topologically associated domains (TADs) during embryogenesis. (A) Adaptions of distribution of radius of gyration of TADs of paternal chromosome. The curves with gray border are the distributions of reference cells. (B) Similar to (A), but for maternal one. (C) Mean radius of gyration of TADs change over time. The lines with gray border represent the values of reference cell. The shadows correspond to the standard deviation of the radius of gyration of TADs divided by 10. (D-F) Similar to (A-C), but for net distance between neighboring TADs. (G) Fractal relationship between net Euclidean distance and sequence distance in the unit of TAD for the case of paternal chromosome. The curves with gray border represent reference cells. (H) Similar to (G), but for maternal one. (I) Fitting parameters of the fractal curves in (G-H). The lines with gray border represent the values of reference cell. The shadows represent the error of fitting (see the supplementary information for details). (J-L) Similar to (G-I), but for those in the unit of 0.1 Mb.

## 3 Conclusions

In this study, we explored the intricate relationship among chromosomal structure, dynamics, and function during embryogenesis using a landscape-switching model based on coarse-grained molecular dynamics simulations. Our model successfully generates continuous ensembles of chromosomal structures that bridge two distinct cell states, thereby predicting the intermediate configurations by which a chromosome transitions from one state to another, a process that is challenging to capture experimentally due to limited temporal resolution. Importantly, the intermediate states predicted by our model align well with experimental observations, underscoring its potential to reveal the underlying mechanisms governing chromosomal reorganization during embryogenesis.

Our analyses reveal that both allelic chromosomes exhibit symmetric developmental behaviors, either both monotonic or both non-monotonic, in terms of higherorder chromosomal structures, chromosomal loci distribution, and short-range TAD organization (Figure 6). Notably, the re-localization of chromosomal loci, particularly gene regions, is decoupled from ZGA and occurs much later, coinciding with the maturation of repressive epigenetic marks that down-regulate totipotency-associated genes and promote the transition to pluripotency. This suggests that the primary functional consequence of gene re-localization is to reposition compartment B inward, thereby repressing specific gene sets, while the outward movement of compartment A appears to be a secondary effect. Consequently, the chromosomal reorganization associated with ZGA likely focuses on TAD dynamics, wherein TADs compact internally to enhance local enhancer-promoter interactions and extend inter-TAD distances to prevent aberrant contacts, thereby establishing a robust gene regulatory network early in embryogenesis.

**Figure 6:**
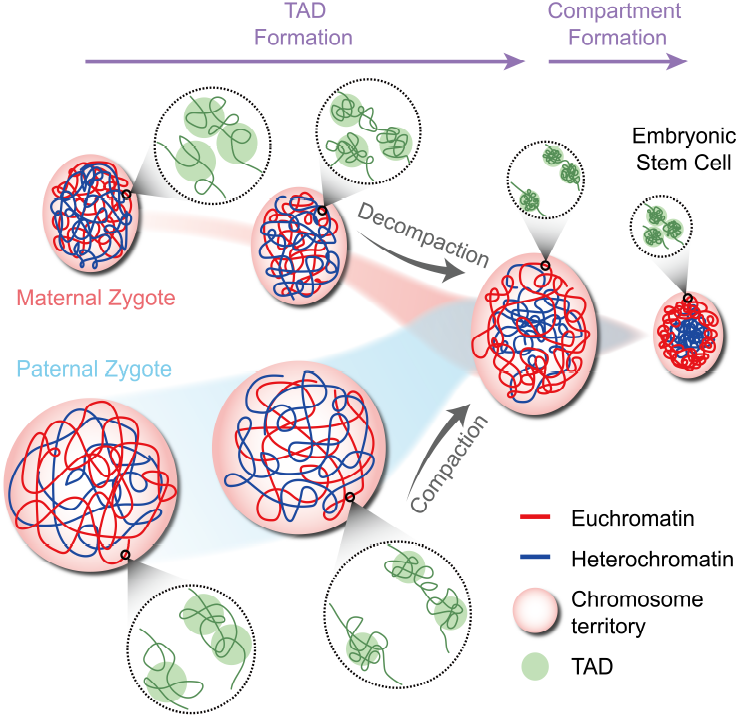
The scheme illustrating of chromosome structure reorganization during embryogenesis. The two leftmost chromosomes are those of ZP and ZM respectively. The rightmost one is that in ESC. The two middle columns are those at intermediate stages. The two allelic development pathways merge at around the third column. The pink globules are the outlines of chromosomes, or equivalent to chromosome territories. The zoomed in images exhibit TADs. Each TAD is outlined by a green globule.

We further identified distinct asymmetric behaviors in chromosome geometry and long-range organization (Figure 6). Specifically, the maternal chromosome exhibits a pronounced over-expansion during embryogenesis, a feature largely driven by changes in its long-range structure, whereas the paternal chromosome undergoes a steady, monotonic change. Although such over-expansion is generally attributed to global DNA demethylation, which relaxes chromatin organization, current evidence does not support a significant difference in DNA demethylation levels between the two allelic chromosomes. Instead, allele-specific histone modifications may better account for these observations. In particular, repressive marks such as H3K27me3, H3K9me3, and H3K64me3, known to promote chromatin compaction, are suggested to be less abundant in the maternal genome, thereby contributing to its over-expanded state during embryogenesis.

In summary, our integrated modeling framework not only recapitulates key experimental observations but also unveils novel insights into the mechanisms underlying chromosomal reorganization during embryogenesis. By bridging early zygotic states and later pluripotent configurations through continuous chromosome structural trajectories, our study highlights the interplay between TAD dynamics, compartment reorganization, and allele-specific behaviors. These findings provide a robust theoretical foundation for understanding how chromosomal architecture regulates gene expression during early development, and they open new avenues for experimental validation and potential applications in regenerative medicine and congenital disease treatment.

## 4 Materials and Methods

### 4.1 Processing of Experimental Data

Hi-C datasets used in this study were obtained from the Gene Expression Omnibus (GEO) under accession number GSE82185 ^[19]^. Raw sequencing reads were aligned to the mouse reference genome (mm9) and subsequently processed and iteratively corrected ^[92]^ using the standard HiC-Pro (version 3.0.0) pipeline ^[93]^. Our analysis focused on the 25–85 Mb region of chromosome 15, with the Hi-C data binned at a resolution of 100 kb. This bin size was chosen to ensure the resulted polymer model, with each bead representing one bin, physically corresponds to the 30-nm chromatin fiber, as noted in previous studies ^[52, 94]^.

To facilitate the calculation of contacts in simulations, the raw contact frequency *f* in Hi-C was normalized to obtain the contact probability *P* under the assumption that adjacent beads are always in contact, i.e. *P*_*i*,*i*±1_≡ 1. This normalization was achieved using the following formula ^[51]^:

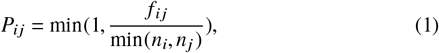

where

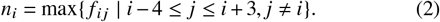

Additionally, gene location data corresponding to the analyzed chromosomal segment were processed using R (version 4.4) in conjunction with Bioconductor (version 3.20). The gene coordinates were binned according to the same 100 kb intervals and subsequently normalized by the total gene count to yield a normalized gene density profile

### 4.2 Chromosome Model

#### 4.2.1 Polymer Model

In our study, the chromosome is represented as a beads-on-a-string homopolymer, which captures key physical characteristics such as chain connectivity, local stiffness, and overall flexibility, etc ^[42, 51, 95]^. Each bead corresponds to a defined chromatin segment, and the system’s dynamics are governed by the total potential energy:

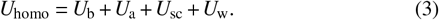

Here, *U*_b_ ensures proper bond connectivity along the chain, while *U*_a_ enforces bending rigidity by penalizing deviations in bond angles, thereby modeling the stiffness of the chromatin fiber ^[95]^. The soft-core repulsive potential *U*_sc_ acts on all non-bonded bead pairs to penalize excessive overlap; this term also permits chain crossing events that mimic the enzymatic activity of topoisomerases, which alleviate topological constraints ^[96]^. Finally, *U*_w_ is a spherical confinement potential that replicates the effects of the nuclear envelope, maintaining the chromosome within a defined volume ^[42, 51, 95]^. Details can be found in supplementary information.

#### 4.2.2 Data-driven 3D Structure Model Based on the Maximum Entropy Principle

To capture the sequence-specific details of chromosomal organization beyond the generic polymer properties, we integrated experimental Hi-C data into our model via the maximum entropy principle ^[51, 52, 97]^. In this framework, the total potential energy *U* is expressed as the sum of the baseline homopolymer energy *U*_homo_ and an additional biasing potential *U*_Hi−C_, which is formulated as a linear combination of observables that reflect contact probabilities between chromosomal loci:

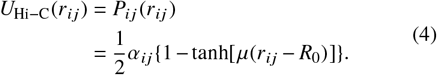

where*r*_*ij*_ denotes the distance between beads *i* and *j*. The ensemble average of the function *P*_*ij*_ (*r*_*ij*_ ) corresponds to the contact probability *P*_*ij*_ obtained from Hi-C data. Thus, the connection between potential energy and observables is built. For the parameters, the *R*_0_ and *µ* are set to 1.5*σ* and 1.2*σ*^−1^, respectively, with *σ* representing the bond length. The coefficients *α*_*ij*_ are iteratively optimized such that the ensemble-averaged contact probabilities computed from the simulated chromosome structures converge to the experimentally derived Hi-C contact probabilities ^[51, 52, 97]^ (Figure S1-S3). Due to the local interactions (*U*_b_ and *U*_a_) span at most 3 consecutive beads, the non-local interactions (*U*_sc_ and *U*_Hi−C_) take effect from beads separated by 3 bonds.

#### 4.2.3 Physics-based Landscape-switching Model

To capture the dynamic transitions between distinct cellular states, we employ a physics-based landscape-switching model ^[42–49]^ that extends our maximum entropy framework. For each transition, from the ZP or ZM state to the ESC state, separate maximum entropy models are generated based on the corresponding Hi-C data (Figure S4). Initially, the chromosome is equilibrated within the energy landscape characteristic of its starting state (either ZP or ZM). At the onset of the cell-state transition, this landscape is instantaneously switched to that of the ESC state. Because the chromosomal structure does not adjust immediately, it gradually relaxes toward the new equilibrium defined by the ESC energy landscape. Such a relaxation process is regarded as the chromosome state transition process. Multiple parallel simulations are performed to generate a series of transition trajectories. At each time point, the ensemble of chromosome structures represents a quasi-equilibrium state, thereby providing a continuous pathway that bridges the initial and final chromosomal configurations.

### 4.3 MD Simulations

All molecular dynamics simulations were conducted using GROMACS ^[98]^ (version 4.5.7) in conjunction with PLUMED ^[99]^ (version 2.5.0), which was employed to implement the spherical confinement potential *U*_w_. A time integration step of 0.0005*τ* was used, where the *τ* denotes the reduced time unit. Langevin dynamics was applied throughout, with a friction coefficient set to 10*τ*^−1^ to ensure proper thermal equilibration. The system temperature was maintained at 1*ϵ* , where *ϵ* represents the intrinsic energy scale; this parameter serves as a relative measure of thermal fluctuations rather than an absolute temperature. All simulations were performed under these standardized conditions unless noted otherwise.

## Acknowledgments

C.F. thanks Tao Zhu and Xingyue Guan for providing technical guidance on the establishment of the maximum entropy model, and Dr. Zhenhai Du for providing technical guidance on the processing of allele-specific Hi-C data. X.C. acknowledges support from the National Natural Science Foundation of China (Grant Nos. 32201020 and 12474201), the general program of Guangdong Basic and Applied Basic Research Foundation (Grant No. 2024A1515010862) and Guangdong provincial project (Grant No. 2023QN10X037).

## References

[1] Li-quan Zhou and Jurrien Dean. Reprogramming the genome to totipotency in mouse embryos. Trends in cell biology, 25(2):82–91, 2015.

[2] Weikun Xia and Wei Xie. Rebooting the epigenomes during mammalian early embryogenesis. Stem cell reports, 15(6):1158–1175, 2020.

[3] Qianhua Xu and Wei Xie. Epigenome in early mammalian development: inheritance, reprogramming and establishment. Trends in cell biology, 28(3):237–253, 2018.

[4] Adam Burton and Maria-Elena Torres-Padilla. Chromatin dynamics in the regulation of cell fate allocation during early embryogenesis. Nature reviews Molecular cell biology, 15(11):723–735, 2014.

[5] Zhenhai Du, Ke Zhang, and Wei Xie. Epigenetic reprogramming in early animal development. Cold Spring Harbor Perspectives in Biology, 14(6):a039677, 2022.

[6] Franka J Rang, Jop Kind, and Isabel Guerreiro. The role of heterochromatin in 3d genome organization during preimplantation development. Cell Reports, 42(4), 2023.

[7] Antoine Vallot and Kikuë Tachibana. The emergence of genome architecture and zygotic genome activation. Current Opinion in Cell Biology, 64:50–57, 2020.

[8] Natasha Jansz and Maria-Elena Torres-Padilla. Genome activation and architecture in the early mammalian embryo. Current opinion in genetics & development, 55:52–58, 2019.

[9] Elizabeth Ing-Simmons, Maria Rigau, and Juan M Vaquerizas. Emerging mechanisms and dynamics of three-dimensional genome organisation at zygotic genome activation. Current Opinion in Cell Biology, 74:37–46, 2022.

[10] Yu Zhang and Wei Xie. Building the genome architecture during the maternal to zygotic transition. Current Opinion in Genetics & Development, 72:91–100, 2022.

[11] Job Dekker, Karsten Rippe, Martijn Dekker, and Nancy Kleckner. Capturing chromosome conformation. science, 295(5558):1306–1311, 2002.

[12] Erez Lieberman-Aiden, Nynke L Van Berkum, Louise Williams, Maxim Imakaev, Tobias Ragoczy, Agnes Telling, Ido Amit, Bryan R Lajoie, Peter J Sabo, Michael O Dorschner, et al. Comprehensive mapping of long-range interactions reveals folding principles of the human genome. science, 326(5950):289–293, 2009.

[13] Guang Shi and Dave Thirumalai. Conformational heterogeneity in human interphase chromosome organization reconciles the fish and hi-c paradox. Nature communications, 10(1):3894, 2019.

[14] Jesse R Dixon, Siddarth Selvaraj, Feng Yue, Audrey Kim, Yan Li, Yin Shen, Ming Hu, Jun S Liu, and Bing Ren. Topological domains in mammalian genomes identified by analysis of chromatin interactions. Nature, 485(7398):376–380, 2012.

[15] Elphège P Nora, Bryan R Lajoie, Edda G Schulz, Luca Giorgetti, Ikuhiro Okamoto, Nicolas Servant, Tristan Piolot, Nynke L Van Berkum, Johannes Meisig, John Sedat, et al. Spatial partitioning of the regulatory landscape of the x-inactivation centre. Nature, 485(7398):381–385, 2012.

[16] Tom Sexton, Eitan Yaffe, Ephraim Kenigsberg, Frédéric Bantignies, Benjamin Leblanc, Michael Hoichman, Hugues Parrinello, Amos Tanay, and Giacomo Cavalli. Three-dimensional folding and functional organization principles of the drosophila genome. Cell, 148(3):458–472, 2012.

[17] Suhas SP Rao, Miriam H Huntley, Neva C Durand, Elena K Stamenova, Ivan D Bochkov, James T Robinson, Adrian L Sanborn, Ido Machol, Arina D Omer, Eric S Lander, et al. A 3d map of the human genome at kilobase resolution reveals principles of chromatin looping. Cell, 159(7):1665–1680, 2014.

[18] Anette Kurz, Stefan Lampel, Jeremy E Nickolenko, Joachim Bradl, Axel Benner, Rebekka M Zirbel, Thomas Cremer, and Peter Lichter. Active and inactive genes localize preferentially in the periphery of chromosome territories. The Journal of cell biology, 135(5):1195–1205, 1996.

[19] Zhenhai Du, Hui Zheng, Bo Huang, Rui Ma, Jingyi Wu, Xianglin Zhang, Jing He, Yunlong Xiang, Qiujun Wang, Yuanyuan Li, et al. Allelic reprogramming of 3d chromatin architecture during early mammalian development. Nature, 547(7662):232–235, 2017.

[20] Samuel Collombet, Noémie Ranisavljevic, Takashi Nagano, Csilla Varnai, Tarak Shisode, Wing Leung, Tristan Piolot, Rafael Galupa, Maud Borensztein, Nicolas Servant, et al. Parental-to-embryo switch of chromosome organization in early embryogenesis. Nature, 580(7801):142– 146, 2020.

[21] Yuwen Ke, Yanan Xu, Xuepeng Chen, Songjie Feng, Zhenbo Liu, Yaoyu Sun, Xuelong Yao, Fangzhen Li, Wei Zhu, Lei Gao, et al. 3d chromatin structures of mature gametes and structural reprogramming during mammalian embryogenesis. Cell, 170(2):367–381, 2017.

[22] Jingyi Wu, BO Huang, HE Chen, Qiangzong Yin, Yang Liu, Yunlong Xiang, Bingjie Zhang, Bofeng Liu, Qiujun Wang, Weikun Xia, et al. The landscape of accessible chromatin in mammalian preimplantation embryos. Nature, 534(7609):652–657, 2016.

[23] Falong Lu, Yuting Liu, Azusa Inoue, Tsukasa Suzuki, Keji Zhao, and Yi Zhang. Establishing chromatin regulatory landscape during mouse preimplantation development. Cell, 165(6):1375–1388, 2016.

[24] Timothy G Jenkins and Douglas T Carrell. Dynamic alterations in the paternal epigenetic landscape following fertilization. Frontiers in genetics, 3:143, 2012.

[25] Richard M Schultz. The molecular foundations of the maternal to zygotic transition in the preimplantation embryo. Human reproduction update, 8(4):323–331, 2002.

[26] Claudia B Walser and Howard D Lipshitz. Transcript clearance during the maternal-to-zygotic transition. Current opinion in genetics & development, 21(4):431–443, 2011.

[27] Petr Svoboda, Vedran Franke, and Richard M Schultz. Sculpting the transcriptome during the oocyte-to-embryo transition in mouse. Current topics in developmental biology, 113:305–349, 2015.

[28] Miler T Lee, Ashley R Bonneau, and Antonio J Giraldez. Zygotic genome activation during the maternal-to-zygotic transition. Annual review of cell and developmental biology, 30(1):581–613, 2014.

[29] Christine Bouniol, Eric Nguyen, and Pascale Debey. Endogenous transcription occurs at the 1-cell stage in the mouse embryo. Experimental cell research, 218(1):57–62, 1995.

[30] JY Nothias, M Miranda, and ML DePamphilis. Uncoupling of transcription and translation during zygotic gene activation in the mouse. The EMBO journal, 15(20):5715–5725, 1996.

[31] Fugaku Aoki, Diane M Worrad, and Richard M Schultz. Regulation of transcriptional activity during the first and second cell cycles in the preimplantation mouse embryo. Developmental biology, 181(2):296– 307, 1997.

[32] Toshio Hamatani, Mark G Carter, Alexei A Sharov, and Minoru SH Ko. Dynamics of global gene expression changes during mouse preimplantation development. Developmental cell, 6(1):117–131, 2004.

[33] Q Tian Wang, Karolina Piotrowska, Maria Anna Ciemerych, Ljiljana Milenkovic, Matthew P Scott, Ronald W Davis, and Magdalena Zernicka-Goetz. A genome-wide study of gene activity reveals developmental signaling pathways in the preimplantation mouse embryo. Developmental cell, 6(1):133–144, 2004.

[34] Fanyi Zeng, Don A Baldwin, and Richard M Schultz. Transcript profiling during preimplantation mouse development. Developmental biology, 272(2):483–496, 2004.

[35] Ken-ichiro Abe, Ryoma Yamamoto, Vedran Franke, Minjun Cao, Yutaka Suzuki, Masataka G Suzuki, Kristian Vlahovicek, Petr Svoboda, Richard M Schultz, and Fugaku Aoki. The first murine zygotic transcription is promiscuous and uncoupled from splicing and 3′ processing. The EMBO journal, 34(11):1523–1537, 2015.

[36] Takashi Nagano, Yaniv Lubling, Tim J Stevens, Stefan Schoenfelder, Eitan Yaffe, Wendy Dean, Ernest D Laue, Amos Tanay, and Peter Fraser. Single-cell hi-c reveals cell-to-cell variability in chromosome structure. Nature, 502(7469):59–64, 2013.

[37] Yusuke Ohnishi, Wolfgang Huber, Akiko Tsumura, Minjung Kang, Panagiotis Xenopoulos, Kazuki Kurimoto, Andrzej K Oleś, Marcos J Araúzo-Bravo, Mitinori Saitou, Anna-Katerina Hadjantonakis, et al. Cell-to-cell expression variability followed by signal reinforcement progressively segregates early mouse lineages. Nature cell biology, 16(1):27–37, 2014.

[38] Arjun Raj and Alexander Van Oudenaarden. Nature, nurture, or chance: stochastic gene expression and its consequences. Cell, 135(2):216–226, 2008.

[39] Gašper Tkačik and William Bialek. Information processing in living systems. Annual Review of Condensed Matter Physics, 7(1):89–117, 2016.

[40] Wendy A Bickmore. The spatial organization of the human genome. Annual review of genomics and human genetics, 14(1):67–84, 2013.

[41] Tom Misteli. Beyond the sequence: cellular organization of genome function. Cell, 128(4):787–800, 2007.

[42] Xiakun Chu and Jin Wang. Conformational state switching and pathways of chromosome dynamics in cell cycle. Applied Physics Reviews, 7(3), 2020.

[43] Xiakun Chu and Jin Wang. Microscopic chromosomal structural and dynamical origin of cell differentiation and reprogramming. Advanced Science, 7(20):2001572, 2020.

[44] Xiakun Chu and Jin Wang. Deciphering the molecular mechanism of the cancer formation by chromosome structural dynamics. PLoS Computational Biology, 17(11):e1009596, 2021.

[45] Xiakun Chu and Jin Wang. Dynamics and pathways of chromosome structural organizations during cell transdifferentiation. JACS Au, 2(1):116–127, 2021.

[46] Xiakun Chu and Jin Wang. Insights into the cell fate decision-making processes from chromosome structural reorganizations. Biophysics Reviews, 3(4), 2022.

[47] Xiakun Chu and Jin Wang. Quantifying chromosome structural reorganizations during differentiation, reprogramming, and transdifferentiation. Physical Review Letters, 129(6):068102, 2022.

[48] Xiakun Chu and Jin Wang. Quantifying the large-scale chromosome structural dynamics during the mitosis-to-g1 phase transition of cell cycle. Open Biology, 13(11):230175, 2023.

[49] Xiakun Chu, Cibo Feng, and Jin Wang. Deciphering the dynamical chromosome structural reorganizations in human neural development. Physical Review Research, 6(2):023309, 2024.

[50] Cibo Feng, Jin Wang, and Xiakun Chu. Large-scale data-driven and physics-based models offer insights into the relationships among the structures, dynamics, and functions of chromosomes. Journal of Molecular Cell Biology, 15(6):mjad042, 2023.

[51] Bin Zhang and Peter G Wolynes. Topology, structures, and energy landscapes of human chromosomes. Proceedings of the National Academy of Sciences, 112(19):6062–6067, 2015.

[52] Bin Zhang and Peter G Wolynes. Shape transitions and chiral symmetry breaking in the energy landscape of the mitotic chromosome. Physical review letters, 116(24):248101, 2016.

[53] Bin Zhang and Peter G Wolynes. Genomic energy landscapes. Biophysical journal, 112(3):427–433, 2017.

[54] Kurt Kremer, Gary S Grest, and I Carmesin. Crossover from rouse to reptation dynamics: A molecular-dynamics simulation. Physical review letters, 61(5):566, 1988.

[55] Jagannathan T Kalathi, Sanat K Kumar, Michael Rubinstein, and Gary S Grest. Rouse mode analysis of chain relaxation in homopolymer melts. Macromolecules, 47(19):6925–6931, 2014.

[56] Ilya M Flyamer, Johanna Gassler, Maxim Imakaev, Hugo B Brandão, Sergey V Ulianov, Nezar Abdennur, Sergey V Razin, Leonid A Mirny, and Kikuë Tachibana-Konwalski. Single-nucleus hi-c reveals unique chromatin reorganization at oocyte-to-zygote transition. Nature, 544(7648):110–114, 2017.

[57] Johanna Gassler, Hugo B Brandão, Maxim Imakaev, Ilya M Flyamer, Sabrina Ladstätter, Wendy A Bickmore, Jan-Michael Peters, Leonid A Mirny, and Kikuë Tachibana. A mechanism of cohesin-dependent loop extrusion organizes zygotic genome architecture. The EMBO journal, 36(24):3600–3618, 2017.

[58] Kristin Abramo, Anne-Laure Valton, Sergey V Venev, Hakan Ozadam, A Nicole Fox, and Job Dekker. A chromosome folding intermediate at the condensin-to-cohesin transition during telophase. Nature cell biology, 21(11):1393–1402, 2019.

[59] Jesse R Dixon, Inkyung Jung, Siddarth Selvaraj, Yin Shen, Jessica E Antosiewicz-Bourget, Ah Young Lee, Zhen Ye, Audrey Kim, Nisha Rajagopal, Wei Xie, et al. Chromatin architecture reorganization during stem cell differentiation. Nature, 518(7539):331–336, 2015.

[60] Emily Crane, Qian Bian, Rachel Patton McCord, Bryan R Lajoie, Bayly S Wheeler, Edward J Ralston, Satoru Uzawa, Job Dekker, and Barbara J Meyer. Condensin-driven remodelling of x chromosome topology during dosage compensation. Nature, 523(7559):240–244, 2015.

[61] Wibke Schwarzer, Nezar Abdennur, Anton Goloborodko, Aleksandra Pekowska, Geoffrey Fudenberg, Yann Loe-Mie, Nuno A Fonseca, Wolfgang Huber, Christian H Haering, Leonid Mirny, et al. Two independent modes of chromatin organization revealed by cohesin removal. Nature, 551(7678):51–56, 2017.

[62] Elphège P Nora, Anton Goloborodko, Anne-Laure Valton, Johan H Gibcus, Alec Uebersohn, Nezar Abdennur, Job Dekker, Leonid A Mirny, and Benoit G Bruneau. Targeted degradation of ctcf decouples local insulation of chromosome domains from genomic compartmentalization. Cell, 169(5):930–944, 2017.

[63] Suhas SP Rao, Su-Chen Huang, Brian Glenn St Hilaire, Jesse M Engreitz, Elizabeth M Perez, Kyong-Rim Kieffer-Kwon, Adrian L Sanborn, Sarah E Johnstone, Gavin D Bascom, Ivan D Bochkov, et al. Cohesin loss eliminates all loop domains. Cell, 171(2):305–320, 2017.

[64] Leonid A Mirny. The fractal globule as a model of chromatin architecture in the cell. Chromosome research, 19:37–51, 2011.

[65] Geoffrey Fudenberg, Maxim Imakaev, Carolyn Lu, Anton Goloborodko, Nezar Abdennur, and Leonid A Mirny. Formation of chromosomal domains by loop extrusion. Cell reports, 15(9):2038–2049, 2016.

[66] Feifei Li, Danyang Wang, Ruigao Song, Chunwei Cao, Zhihua Zhang, Yu Wang, Xiaoli Li, Jiaojiao Huang, Qiang Liu, Naipeng Hou, et al. The asynchronous establishment of chromatin 3d architecture between in vitro fertilized and uniparental preimplantation pig embryos. Genome Biology, 21:1–21, 2020.

[67] Susana Gonzalo, Isabel Jaco, Mario F Fraga, Taiping Chen, En Li, Manel Esteller, and María A Blasco. Dna methyltransferases control telomere length and telomere recombination in mammalian cells. Nature cell biology, 8(4):416–424, 2006.

[68] Jiameng Dan, Philippe Rousseau, Swanand Hardikar, Nicolas Veland, Jiemin Wong, Chantal Autexier, and Taiping Chen. Zscan4 inhibits maintenance dna methylation to facilitate telomere elongation in mouse embryonic stem cells. Cell reports, 20(8):1936–1949, 2017.

[69] Zachary D Smith, Jiantao Shi, Hongcang Gu, Julie Donaghey, Kendell Clement, Davide Cacchiarelli, Andreas Gnirke, Franziska Michor, and Alexander Meissner. Epigenetic restriction of extraembryonic lineages mirrors the somatic transition to cancer. Nature, 549(7673):543–547, 2017.

[70] Yu Zhang, Yunlong Xiang, Qiangzong Yin, Zhenhai Du, Xu Peng, Qiujun Wang, Miguel Fidalgo, Weikun Xia, Yuanyuan Li, Zhen-ao Zhao, et al. Dynamic epigenomic landscapes during early lineage specification in mouse embryos. Nature genetics, 50(1):96–105, 2018.

[71] Hui Zheng, Bo Huang, Bingjie Zhang, Yunlong Xiang, Zhenhai Du, Qianhua Xu, Yuanyuan Li, Qiujun Wang, Jing Ma, Xu Peng, et al. Resetting epigenetic memory by reprogramming of histone modifications in mammals. Molecular cell, 63(6):1066–1079, 2016.

[72] Chenfei Wang, Xiaoyu Liu, Yawei Gao, Lei Yang, Chong Li, Wenqiang Liu, Chuan Chen, Xiaochen Kou, Yanhong Zhao, Jiayu Chen, et al. Reprogramming of h3k9me3-dependent heterochromatin during mammalian embryo development. Nature cell biology, 20(5):620–631, 2018.

[73] Sylvain Daujat, Thomas Weiss, Fabio Mohn, Ulrike C Lange, Céline Ziegler-Birling, Ulrike Zeissler, Michael Lappe, Dirk Schübeler, Maria-Elena Torres-Padilla, and Robert Schneider. H3k64 trimethylation marks heterochromatin and is dynamically remodeled during developmental reprogramming. Nature structural & molecular biology, 16(7):777–781, 2009.

[74] Takumi Takizawa, Karen J Meaburn, and Tom Misteli. The meaning of gene positioning. Cell, 135(1):9–13, 2008.

[75] Ana Pombo and Niall Dillon. Three-dimensional genome architecture: players and mechanisms. Nature reviews Molecular cell biology, 16(4):245–257, 2015.

[76] Fides Zenk, Yinxiu Zhan, Pavel Kos, Eva Löser, Nazerke Atinbayeva, Melanie Schächtle, Guido Tiana, Luca Giorgetti, and Nicola Iovino. Hp1 drives de novo 3d genome reorganization in early drosophila embryos. Nature, 593(7858):289–293, 2021.

[77] Todd S Macfarlan, Wesley D Gifford, Saurabh Agarwal, Shawn Driscoll, Karen Lettieri, Jianxun Wang, Shane E Andrews, Laura Franco, Michael G Rosenfeld, Bing Ren, et al. Endogenous retroviruses and neighboring genes are coordinately repressed by lsd1/kdm1a. Genes & development, 25(6):594–607, 2011.

[78] Jamie A Hackett and M Azim Surani. Regulatory principles of pluripotency: from the ground state up. Cell stem cell, 15(4):416–430, 2014.

[79] Máté Borsos and Maria-Elena Torres-Padilla. Building up the nucleus: nuclear organization in the establishment of totipotency and pluripotency during mammalian development. Genes & development, 30(6):611–621, 2016.

[80] Kaixin Wu, HE Liu, Yaofeng Wang, Jiangping He, Shuyang Xu, Yaping Chen, Junqi Kuang, Jiadong Liu, Lin Guo, Dongwei Li, et al. Setdb1-mediated cell fate transition between 2c-like and pluripotent states. Cell reports, 30(1):25–36, 2020.

[81] Yezhang Zhu, Jiali Yu, Jiahui Gu, Chaoran Xue, Long Zhang, Jiekai Chen, and Li Shen. Relaxed 3d genome conformation facilitates the pluripotent to totipotent-like state transition in embryonic stem cells. Nucleic acids research, 49(21):12167–12177, 2021.

[82] Maria Vega-Sendino and Sergio Ruiz. Transition from totipotency to pluripotency in mice: insights into molecular mechanisms. Biochemical Society Transactions, 2024.

[83] Fan Zhou, Rui Wang, Peng Yuan, Yixin Ren, Yunuo Mao, Rong Li, Ying Lian, Junsheng Li, Lu Wen, Liying Yan, et al. Reconstituting the transcriptome and dna methylome landscapes of human implantation. Nature, 572(7771):660–664, 2019.

[84] Weikun Xia, Jiawei Xu, Guang Yu, Guidong Yao, Kai Xu, Xueshan Ma, Nan Zhang, Bofeng Liu, Tong Li, Zili Lin, et al. Resetting histone modifications during human parental-to-zygotic transition. Science, 365(6451):353–360, 2019.

[85] Wenhao Zhang, Zhiyuan Chen, Qiangzong Yin, Dan Zhang, Catherine Racowsky, and Yi Zhang. Maternal-biased h3k27me3 correlates with paternal-specific gene expression in the human morula. Genes & development, 33(7-8):382–387, 2019.

[86] Diego Rodriguez-Terrones, Xavier Gaume, Takashi Ishiuchi, Amélie Weiss, Arnaud Kopp, Kai Kruse, Audrey Penning, Juan M Vaquerizas, Laurent Brino, and Maria-Elena Torres-Padilla. A molecular roadmap for the emergence of early-embryonic-like cells in culture. Nature genetics, 50(1):106–119, 2018.

[87] Ke Zhang, Dan-Ya Wu, Hui Zheng, Yao Wang, Qiao-Ran Sun, Xin Liu, Li-Yan Wang, Wen-Jing Xiong, Qiujun Wang, James DP Rhodes, et al. Analysis of genome architecture during scnt reveals a role of cohesin in impeding minor zga. Molecular cell, 79(2):234–250, 2020.

[88] Mo Chen, Qianshu Zhu, Chong Li, Xiaochen Kou, Yanhong Zhao, Yanhe Li, Ruimin Xu, Lei Yang, Lingyue Yang, Liang Gu, et al. Chromatin architecture reorganization in murine somatic cell nuclear transfer embryos. Nature communications, 11(1):1813, 2020.

[89] Teresa Olbrich, Maria Vega-Sendino, Desiree Tillo, Wei Wu, Nicholas Zolnerowich, Raphael Pavani, Andy D Tran, Catherine N Domingo, Mariajose Franco, Marta Markiewicz-Potoczny, et al. Ctcf is a barrier for 2c-like reprogramming. Nature Communications, 12(1):4856, 2021.

[90] Martin Franke, Elisa De la Calle-Mustienes, Ana Neto, María Almuedo-Castillo, Ibai Irastorza-Azcarate, Rafael D Acemel, Juan J Tena, José M Santos-Pereira, and José L Gómez-Skarmeta. Ctcf knockout in zebrafish induces alterations in regulatory landscapes and developmental gene expression. Nature communications, 12(1):5415, 2021.

[91] Maria Jose Andreu, Alba Alvarez-Franco, Marta Portela, Daniel Gimenez-Llorente, Ana Cuadrado, Claudio Badia-Careaga, Maria Tiana, Ana Losada, and Miguel Manzanares. Establishment of 3d chromatin structure after fertilization and the metabolic switch at the morula-to-blastocyst transition require ctcf. Cell reports, 41(3), 2022.

[92] Maxim Imakaev, Geoffrey Fudenberg, Rachel Patton McCord, Natalia Naumova, Anton Goloborodko, Bryan R Lajoie, Job Dekker, and Leonid A Mirny. Iterative correction of hi-c data reveals hallmarks of chromosome organization. Nature methods, 9(10):999–1003, 2012.

[93] Nicolas Servant, Nelle Varoquaux, Bryan R Lajoie, Eric Viara, Chong-Jian Chen, Jean-Philippe Vert, Edith Heard, Job Dekker, and Emmanuel Barillot. Hic-pro: an optimized and flexible pipeline for hi-c data processing. Genome biology, 16:1–11, 2015.

[94] Fabrizio Benedetti, Julien Dorier, Yannis Burnier, and Andrzej Stasiak. Models that include supercoiling of topological domains reproduce several known features of interphase chromosomes. Nucleic acids research, 42(5):2848–2855, 2014.

[95] Angelo Rosa and Ralf Everaers. Structure and dynamics of interphase chromosomes. PLoS computational biology, 4(8):e1000153, 2008.

[96] Natalia Naumova, Maxim Imakaev, Geoffrey Fudenberg, Ye Zhan, Bryan R Lajoie, Leonid A Mirny, and Job Dekker. Organization of the mitotic chromosome. Science, 342(6161):948–953, 2013.

[97] Andrea Cesari, Sabine Reißer, and Giovanni Bussi. Using the maximum entropy principle to combine simulations and solution experiments. Computation, 6(1):15, 2018.

[98] Berk Hess, Carsten Kutzner, David Van Der Spoel, and Erik Lindahl. Gromacs 4: algorithms for highly efficient, load-balanced, and scalable molecular simulation. Journal of chemical theory and computation, 4(3):435–447, 2008.

[99] Gareth A Tribello, Massimiliano Bonomi, Davide Branduardi, Carlo Camilloni, and Giovanni Bussi. Plumed 2: New feathers for an old bird. Computer physics communications, 185(2):604–613, 2014.

