## Supplementary material for "(A)symmetry in the Allele-Specific Chromosomal Structural Dynamics during Embryogenesis": SI Text

### 1 Supplementary Methods

#### 1.1 Parameterization in the Chromosome Model

##### *1.1.1 Polymer Model*

The bond potential $U_{\mathrm{b}}$ consists of two parts: the finitely extensible nonlinear elastic (FENE) potential $U_{\mathrm{FENE}}$ [1] and the hard-core repulsive potential $U_{\mathrm{hc}}$:

$$U_{\mathrm{b}} = U_{\mathrm{FENE}} + U_{\mathrm{hc}}. \tag{1}$$

The FENE potential

$$U_{\mathrm{FENE}}(r_{i,i+1}) = \begin{cases} -\frac{1}{2}K_{\mathrm{b}}R_0^2 \ln[1-(\frac{r_{i,i+1}}{R_0^2})^2], & \text{if } r_{i,i+1} < R_0^2 - \kappa\sigma \\ K_{\mathrm{p}}(\frac{r_{i,i+1}}{R_0^2})^5, & \text{if } r_{i,i+1} \geq R_0^2 - \kappa\sigma \end{cases} \tag{2}$$

inhibits the breaking of the chain. Here the second term is to avoid the system instability due to the divergence of the first term around $r = R_0$ [2]. The hard-core repulsive potential

$$U_{\mathrm{hc}}(r_{i,i+1}) = \begin{cases} 4\epsilon[(\frac{\sigma}{r_{i,i+1}})^{12} - (\frac{\sigma}{r_{i,i+1}})^6] + \epsilon, & \text{if } r_{i,i+1} < 2^{1/6}\sigma \\ 0, & \text{if } r_{i,i+1} \geq 2^{1/6}\sigma \end{cases} \tag{3}$$

inhibits the overlapping between adjacent beads.

In order to present the stiffness of chromatin fiber [3], the bending potential $U_{\mathrm{a}}$ should make the chain prefer being straight:

$$U_{\mathrm{a}}(\theta_{i-1,i,i+1}) = K_{\mathrm{a}}(1 + \cos\theta_{i-1,i,i+1}). \tag{4}$$

The soft-core repulsive and spherical wall potential are respectively

$$U_{\mathrm{sc}}(r_{ij}) = \begin{cases} \frac{1}{2}E_{\mathrm{cut}}(1 + \tanh\{\frac{8\epsilon}{E_{\mathrm{cut}}}[(\frac{\sigma}{r_{ij}})^{12} - (\frac{\sigma}{r_{ij}})^6] + \frac{8\epsilon}{E_{\mathrm{cut}}} - 1\}), & \text{if } r_{ij} < r_0 \\ 4\epsilon[(\frac{\sigma}{r_{ij}})^{12} - (\frac{\sigma}{r_{ij}})^6] + \epsilon, & \text{if } r_0 \leq r_{ij} < 2^{1/6}\sigma \\ 0, & \text{if } r_{ij} \geq 2^{1/6}\sigma \end{cases}. \tag{5}$$

$$U_{\mathrm{w}}(r) = \begin{cases} 0 & \text{if } r < R_{\mathrm{w}} \\ K_{\mathrm{w}}(r - R_{\mathrm{w}})^2, & \text{if } r \geq R_{\mathrm{w}} \end{cases}. \tag{6}$$

During the simulation, reduced units are used. The energy unit is $\epsilon$. The length is expressed in unit of bond length $\sigma$. Other parameters in the potentials are listed in Table S1.

Table S1:

| Parameters | Values |
|---|---|
| $K_\mathrm{b}$ | $30\epsilon/\sigma$ |
| $R_0$ | $1.5\sigma$ |
| $K_\mathrm{p}$ | $100\epsilon$ |
| $\kappa$ | 0.125 |
| $K_\mathrm{a}$ | $1\epsilon$ |
| $E_\mathrm{cut}$ | $4\epsilon$ |
| $r_0$ | $(\frac{2}{1+\sqrt{2}})^{1/6}\sigma$ |
| $K_\mathrm{w}$ | $100\epsilon/\sigma^2$ |
| $R_\mathrm{w}$ | $13\sigma$ |

##### 1.2 MD Simulation Details

For the iteration of the maximum entropy model, in order to enhance the efficiency and sampling during the stage of performing simulations, 100 independent trajectories are run simultaneously from different initial structures during each epoch. In addition, the simulated annealing technique is employed for each trajectory. In detail, during the first $250\tau$, the temperature is gradually decreased from $4\epsilon$ to $1\epsilon$; from $250\tau$ to $500\tau$, the temperature is constantly $1\epsilon$ for full relaxation; and from $500\tau$ to $1000\tau$, the temperature is also constantly $1\epsilon$ for sampling. The frames are dumped every $0.1\tau$. The required quantities are calculated using the 100 trajectories of $500\tau$, totally $5\times10^5$ frames, to iterate the $\alpha_{ij}$.

For the landscape-switching model, the initial structures of the independent trajectories are from the ensemble obtained during the last epoch of maximum entropy model. However, $5\times10^5$ trajectories are too resource-consuming and redundant. In order to reduce them and meanwhile ensure the efficiency of the time-course ensembles during transitions, trajectories with similar initial structures should be removed, as they will give similar results. To this end, the initial structure ensemble containing $5\times10^5$ structures should be clustered. Under a proper cut-off, only one representative structure in each cluster are selected and grouped into a new initial ensemble, with broad and proper number of structures to be used to perform the subsequent landscape-switching simulations. The selection of the cut-off is based on the accessibility range of each structure during the equilibration diffusion. For example, the chromosome of paternal zygote can deviate from the initiate structure by at most $d_\mathrm{rms} = 6\sigma$ (Figure S4A). The cut-off of dividing the clusters is thus set as $6\sigma$. The case of maternal one is similar (Figure S4B). This results the new initial ensemble of paternal and maternal zygote with 1213 and 973 structures respectively. In addition, the reduced ensemble can reproduce the average results of the raw ensemble to an extremely large extent (Figure S4E-P), which ensures the rationality of the reduction.

Obtaining the reduced initial structure ensemble, 1213 (973) simulation trajectories for paternal (maternal) development are conducted according to the landscape-switching model. Firstly, the equilibrium simulations under the potential of zygote are run for $5000\tau$, simulating the stable zygote states before embryogenesis beginning. Subsequently, the potentials are suddenly changed, indicating the embryogenesis begins. The simulations under the new potential are run for $1000\tau$, allowing full relaxation mimicking the embryogenesis progressing. The frames during this process are dumped every $0.01\tau$ for the subsequent analysis.

##### 1.3 Calculation Methods of Analysis Quantities

When characterizing the chromosome structures, contact probability maps are widely used, averaging out the ensemble heterogeneity and directly corresponding to the experimental measurements. The contact probability is calculated by

$$P_{ij} = \frac{1}{2}\{1 - \tanh[\mu(r_{ij} - R_0)]\}. \tag{7}$$

In order to investigate the structures based on contact maps, the coefficient of determination [4]

$$R^2(P, P^{\mathrm{ref}}) = 1 - \frac{\sum_{i,j}(P_{ij} - P^{\mathrm{ref}}_{ij})^2}{\sum_{i,j}(P_{ij} - \overline{P})^2}, \tag{8}$$

is used to quantify the similarity between the contact map of chromosome in transition $P$ and that of reference cells $P^{\mathrm{ref}}$, like zygotes, 8-cell states, and ESC. Here $\overline{P}$ is the mean over all pairs $(i, j)$. A higher value of $R^2$ indicates they are more similar, and the maximum $R^2$ equal to 1 means they are identical actually. In addition, to enable more meaningful comparison, the contact probabilities $P$ are subtracted by the distance factors

$$p_\xi = \frac{1}{N-\xi}\sum_{i=1}^{N-\xi} P_{i,i+\xi}, \tag{9}$$

where $\xi$ is the genomic distance and $N$ is the number of all beads. This is because $p$ is the common feature of all polymers regardless of higher-order chromosome structures.

To quantify the compartment, the enhanced contact matrix

$$S_{\mathrm{C}} = \log_2(P^{\mathrm{Obs}}/P^{\mathrm{Exp}}) \tag{10}$$

is calculated [5,6]. Here $P^{\mathrm{Obs}}$ is the coarse-grained contact probability $P$ at a bin size of 1 Mb (10 times original bin size). $P^{\mathrm{Exp}}$ satisfies

$$P^{\mathrm{Exp}}_{i,i+\xi} = p_\xi \ \forall i < N - \xi. \tag{11}$$

In order to calculate the compartmentalization strength $A_{\mathrm{C}}$, the following program [7] is performed. Firstly, the one-dimensional compartment profile, i.e. the first principal component of $S_{\mathrm{C}}$, of ESC is denoted as $S_{\mathrm{c}}^{\mathrm{ESC}}$ and modified according to the gene density profile, making the correlation between them is positive. Secondly, $S_{\mathrm{C}}$ is sorted by $S_{\mathrm{c}}^{\mathrm{ESC}}$ in both dimensions. Thirdly, the entries of sorted $S_{\mathrm{C}}$ in the corner corresponding to positive $S_{\mathrm{c}}^{\mathrm{ESC}}$ in both dimensions are averaged as $\overline{S_{\mathrm{c}}^{\mathrm{A-A}}}$, negative $S_{\mathrm{c}}^{\mathrm{ESC}}$ in both dimensions are averaged as $\overline{S_{\mathrm{c}}^{\mathrm{B-B}}}$, and the remain corners are averaged as $\overline{S_{\mathrm{c}}^{\mathrm{A-B}}}$. Lastly, the $A_{\mathrm{C}}$ is defined as

$$A_{\mathrm{c}} = \frac{\overline{S_{\mathrm{c}}^{\mathrm{A-A}}} + \overline{S_{\mathrm{c}}^{\mathrm{B-B}}}}{\overline{S_{\mathrm{c}}^{\mathrm{A-B}}}}. \tag{12}$$

The insulation score $S_{\mathrm{i}}$ is calculated referring to [8]. The TAD boundaries are defined according to ESC. The insulation strength $A_{\mathrm{i}}$ is defined as follows [7,9]. The $S_{\mathrm{i}}$ around all TAD boundaries by ±0.5 Mb are averaged as $\overline{S_{\mathrm{i}}}$. It gives a V-shape curve. Then the $A_{\mathrm{i}}$ is defined as

$$A_{\mathrm{i}} = \max(\overline{S_{\mathrm{i}}}) - \min(\overline{S_{\mathrm{i}}}). \tag{13}$$

The polymer state is defined as the logarithm slope of $p$ v.s. $\xi$ curve in the $\xi$ range of 0.5-7 Mb:

$$\gamma = \frac{\mathrm{d}\log p}{\mathrm{d}\log \xi} = \frac{\xi \mathrm{d}p}{p \mathrm{d}\xi} \tag{14}$$

It is the power exponent of power law $p \propto \xi^{\gamma}$ fitted from the data. A higher value of $\gamma$ indicates a more compact polymer state.

For the $d$ v.s. $\xi$ curve, the parameter $\omega$ is defined as

$$\omega = \frac{1}{N-100} \sum_{\xi=100}^{N} \frac{d}{\xi^{0.2}}. \tag{15}$$

It is actually also the fitting by $d = \omega \xi^{0.2}$. Because the analysis based on $d$ more focuses on the long-scale structures, we only consider the distance range from 100 to $N$ beads. Conversely, a higher value of $\omega$ indicates a more relaxed conformation. The corresponding error is

$$E_{\omega} = \frac{1}{N-100} \sum_{\xi=100}^{N} (\frac{d}{\xi^{0.2}} - \omega)^2. \tag{16}$$

It measures the reliability of the fitting.

The compactness $P_{\mathbb{L}}$ within the length scale $\mathbb{L} \in \{0-2\text{Mb}, 2-6\text{Mb}, 6-14\text{Mb}, 14-30\text{Mb}\}$ is defined as

$$P_{\mathbb{L}} = \frac{1}{N_{\Sigma}} \sum_{|i-j| \in \mathbb{L}} P_{ij}. \tag{17}$$

It is the mean value of contact probability within the corresponding range. A higher value indicates a more compact structure within this range. The corresponding folding progress is

$$Q_{\mathbb{L}} = \frac{1}{N_{\Sigma}} \sum_{|i-j| \in \mathbb{L}} \exp[\frac{(r_{ij} - r_{ij}^{\text{ESC}})^2}{2(\sigma/2)^2}], \tag{18}$$

which actually quantifies the similarity between the structure in transition and that of ESC, widely used for protein folding. A higher value indicates the structure is closer to the folding destination.

For kinetics, unless mean behavior, we are also concerned about the heterogeneity among loci. The interactions formed by each locus with other loci at different genomic distance can be calculated by

$$P_{i\mathbb{L}} = \frac{1}{N_{\Sigma}} \sum_{\{j \mid |i-j| \in \mathbb{L}\}} P_{ij}, \tag{19}$$

and

$$Q_{i\mathbb{L}} = \frac{1}{N_{\Sigma}} \sum_{\{j \mid |i-j| \in \mathbb{L}\}} \exp[\frac{(r_{ij} - r_{ij}^{\text{ESC}})^2}{2(\sigma/2)^2}]. \tag{20}$$

There are similar to the $P_{\mathbb{L}}$ and $Q_{\mathbb{L}}$, but the averaging is implemented for each locus, rather than all loci. They are temporally normalized by the function

$$\chi(q(t)) = \frac{q(t) - q(t = 10^{-2}\tau)}{q(t = 10^{3}\tau) - q(t = 10^{-2}\tau)}. \tag{21}$$

The half life $T_{1/2}$ is defined as the time when $\chi(P_{i\mathbb{L}})$ or $\chi(Q_{i\mathbb{L}})$ reaches to $\frac{1}{2}$ for the first time.

For the shape parameters, the basic quantity is the inertial tensor

$$I = \begin{pmatrix} x_1 & x_2 & \dots & x_N \\ y_1 & y_2 & \dots & y_N \\ z_1 & z_2 & \dots & z_N \end{pmatrix} \begin{pmatrix} x_1 & y_1 & z_1 \\ x_2 & y_2 & z_2 \\ \dots & \dots & \dots \\ x_N & y_N & z_N \end{pmatrix}. \tag{22}$$

Here $(x_i, y_i, z_i)$ is the Cartesian coordinates of the $i$-th bead. The $i$-th eigenvalue of $I$ are denoted as $\lambda_i$. The inertial principal axis length

$$R_{\mathrm{PA}i} = \sqrt{\lambda_i}, \tag{23}$$

the radius of gyration

$$R_{\mathrm{g}} = \sqrt{\mathrm{tr}I} = \sqrt{\sum_{i=1}^{3} \lambda_i}, \tag{24}$$

the aspherical parameter

$$\varDelta = \frac{3}{2} \frac{\sum_{i=1}^{3} (\lambda_i - \overline{\lambda})^2}{(\mathrm{tr}I)^2}, \tag{25}$$

and

$$S = 27 \frac{\prod_{i=1}^{3} (\lambda_i - \overline{\lambda})}{(\mathrm{tr}I)^3}. \tag{26}$$

The value of $\varDelta$ is in the range between 0 and 1. A larger value indicates more deviation from sphere shape. Positive and negative value of $S$ means tube-like and disk-like shapes respectively.

In order to quantify the shift direction of the $d^{\mathrm{TAD}}$ v.s. $\xi^{\mathrm{TAD}}$ curve, similar to the case of $d$ v.s. $\xi$, the curve parameter $\varOmega$ is defined as

$$\varOmega = \frac{1}{N^{\mathrm{TAD}} - 10} \sum_{\xi^{\mathrm{TAD}}=10}^{N^{\mathrm{TAD}}} \frac{d^{\mathrm{TAD}}}{\ln \xi^{\mathrm{TAD}}}, \tag{27}$$

where $N^{\mathrm{TAD}}$ is the number of TAD in ESC. It is actually also the fitting by $d^{\mathrm{TAD}} = \varOmega \ln \xi^{\mathrm{TAD}}$. Because the involved analysis more focuses on the long-scale inter-TAD structures, we only consider the distance range from 10 to $N^{\mathrm{TAD}}$ TADs. The corresponding error is

$$E_{\varOmega} = \frac{1}{N^{\mathrm{TAD}} - 10} \sum_{\xi^{\mathrm{TAD}}=10}^{N^{\mathrm{TAD}}} \left(\frac{d^{\mathrm{TAD}}}{\ln \xi^{\mathrm{TAD}}} - \varOmega\right)^2. \tag{28}$$

It also measures the reliability of the fitting.

### References


[1] Harold R Warner Jr. Kinetic theory and rheology of dilute suspensions of finitely extendible dumbbells. *Industrial & Engineering Chemistry Fundamentals*, 11(3):379–387, 1972.

[2] Naoko Tokuda, Tomoki P Terada, and Masaki Sasai. Dynamical modeling of three-dimensional genome organization in interphase budding yeast. *Biophysical journal*, 102(2):296–304, 2012.

[3] Angelo Rosa and Ralf Everaers. Structure and dynamics of interphase chromosomes. *PLoS computational biology*, 4(8):e1000153, 2008.

[4] Xiakun Chu, Cibo Feng, and Jin Wang. Deciphering the dynamical chromosome structural reorganizations in human neural development. *Physical Review Research*, 6(2):023309, 2024.

[5] Erez Lieberman-Aiden, Nynke L Van Berkum, Louise Williams, Maxim Imakaev, Tobias Ragoczy, Agnes Telling, Ido Amit, Bryan R Lajoie, Peter J Sabo, Michael O Dorschner, et al. Comprehensive mapping of long-range interactions reveals folding principles of the human genome. *science*, 326(5950):289–293, 2009.

[6] Jesse R Dixon, Inkyung Jung, Siddarth Selvaraj, Yin Shen, Jessica E Antosiewicz-Bourget, Ah Young Lee, Zhen Ye, Audrey Kim, Nisha Rajagopal, Wei Xie, et al. Chromatin architecture reorganization during stem cell differentiation. *Nature*, 518(7539):331–336, 2015.

[7] Kristin Abramo, Anne-Laure Valton, Sergey V Venev, Hakan Ozadam, A Nicole Fox, and Job Dekker. A chromosome folding intermediate at the condensin-to-cohesin transition during telophase. *Nature cell biology*, 21(11):1393–1402, 2019.

[8] Emily Crane, Qian Bian, Rachel Patton McCord, Bryan R Lajoie, Bayly S Wheeler, Edward J Ralston, Satoru Uzawa, Job Dekker, and Barbara J Meyer. Condensin-driven remodelling of x chromosome topology during dosage compensation. *Nature*, 523(7559):240–244, 2015.

[9] Zhenhai Du, Hui Zheng, Bo Huang, Rui Ma, Jingyi Wu, Xianglin Zhang, Jing He, Yunlong Xiang, Qiujun Wang, Yuanyuan Li, et al. Allelic reprogramming of 3d chromatin architecture during early mammalian development. *Nature*, 547(7662):232–235, 2017.

#### 2 Supplementary Figures

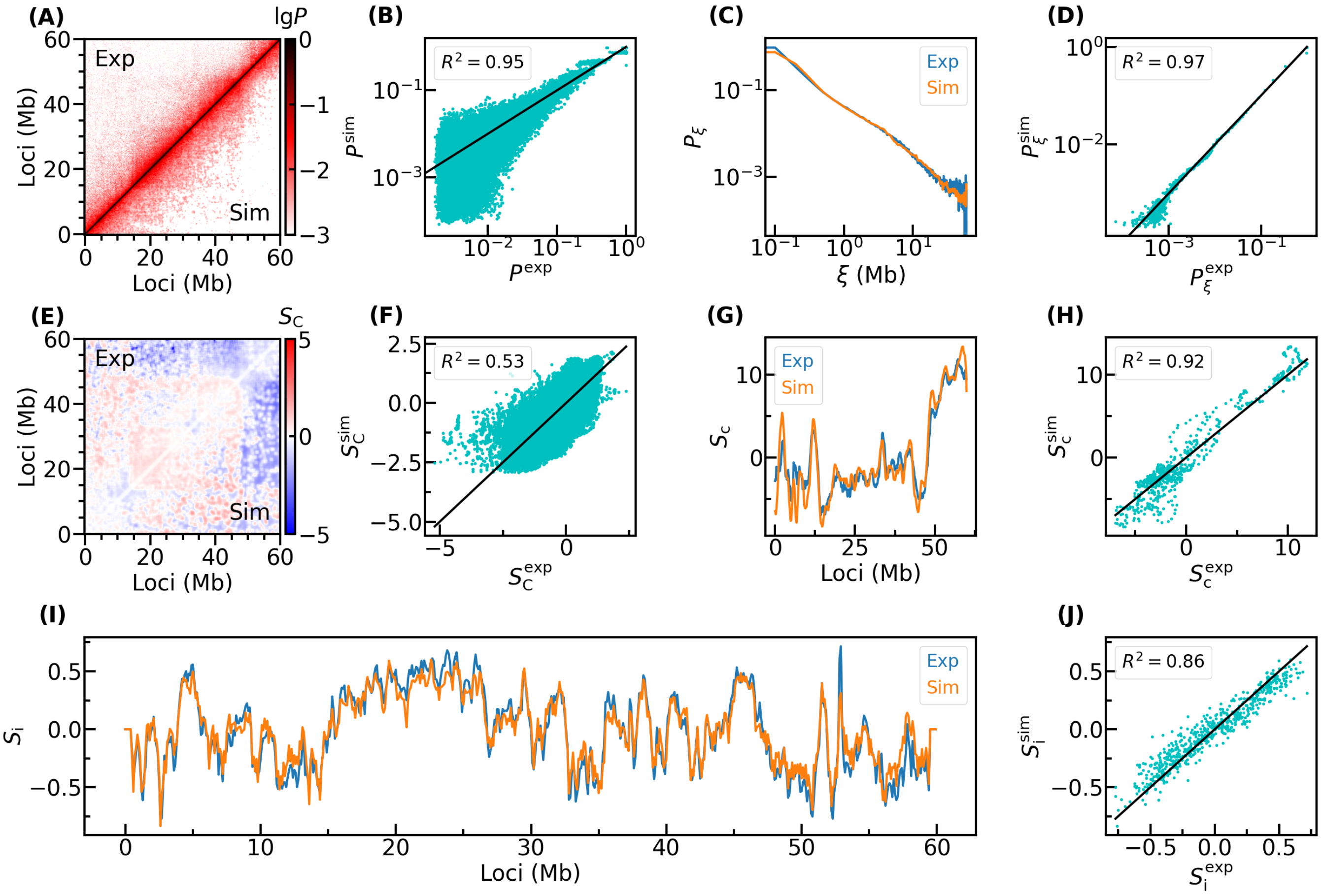


Figure S1: Validation of the maximum entropy model for the chromosome structures of ZP. (A) Comparison between contact probability matrix from Hi-C experiment (the upper left triangle, denoted as $P^{\text{exp}}$) and from maximum entropy model simulation (the lower right triangle, denoted as $P^{\text{sim}}$). (B) Scatter correlation between $P^{\text{exp}}$ and $P^{\text{sim}}$. The black line represents $P^{\text{sim}} = P^{\text{exp}}$. The legend shows the coefficient of determination between $P^{\text{exp}}$ and $P^{\text{sim}}$. (C) Comparison of $P_\xi$ decay curve. (D) Scatter correlation between $P_\xi^{\text{exp}}$ and $P_\xi^{\text{sim}}$, which are the value obtained from experiment and simulation. (E) Comparison of $S_{\text{C}}$ matrix. (F) Corresponding scatter correlation. (G) Comparison of 1D compartment profile. (H) Corresponding scatter correlation. (I-K) Comparison of insulation score, compared in three separate segments. (L) Corresponding scatter correlation.

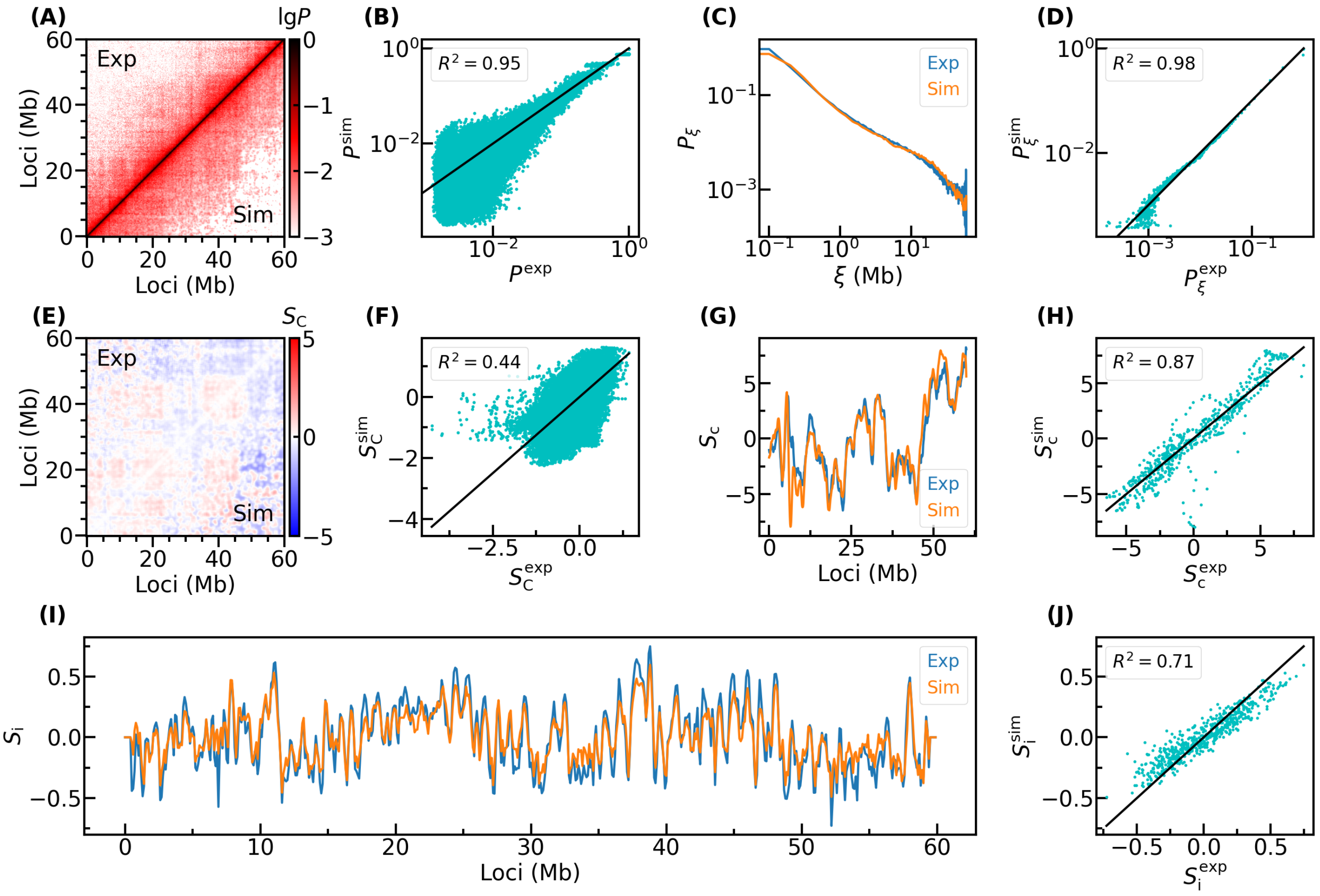


Figure S2: Similar to Figure S1, but for ZM.

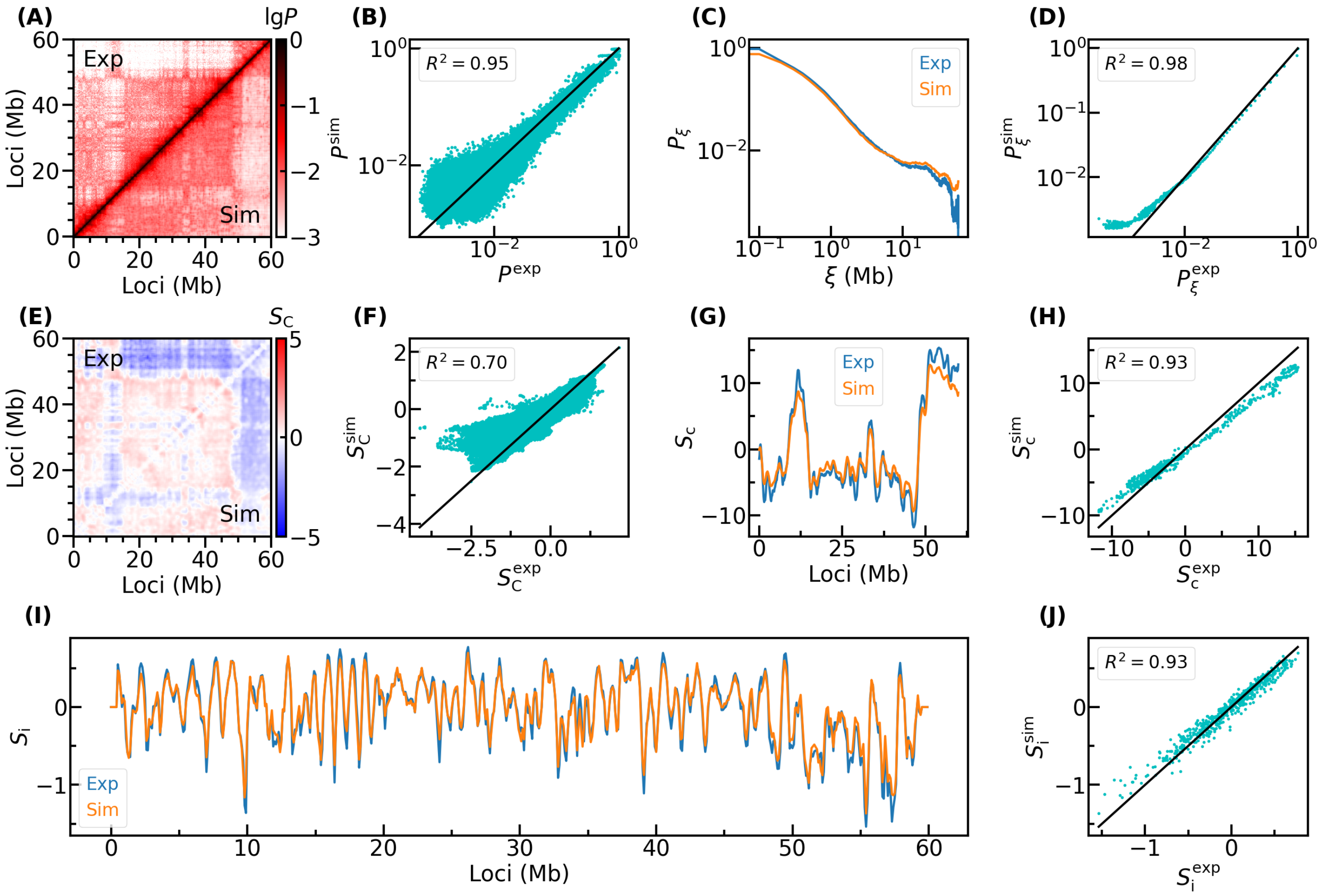


Figure S3: Similar to Figure S1, but for ESC.

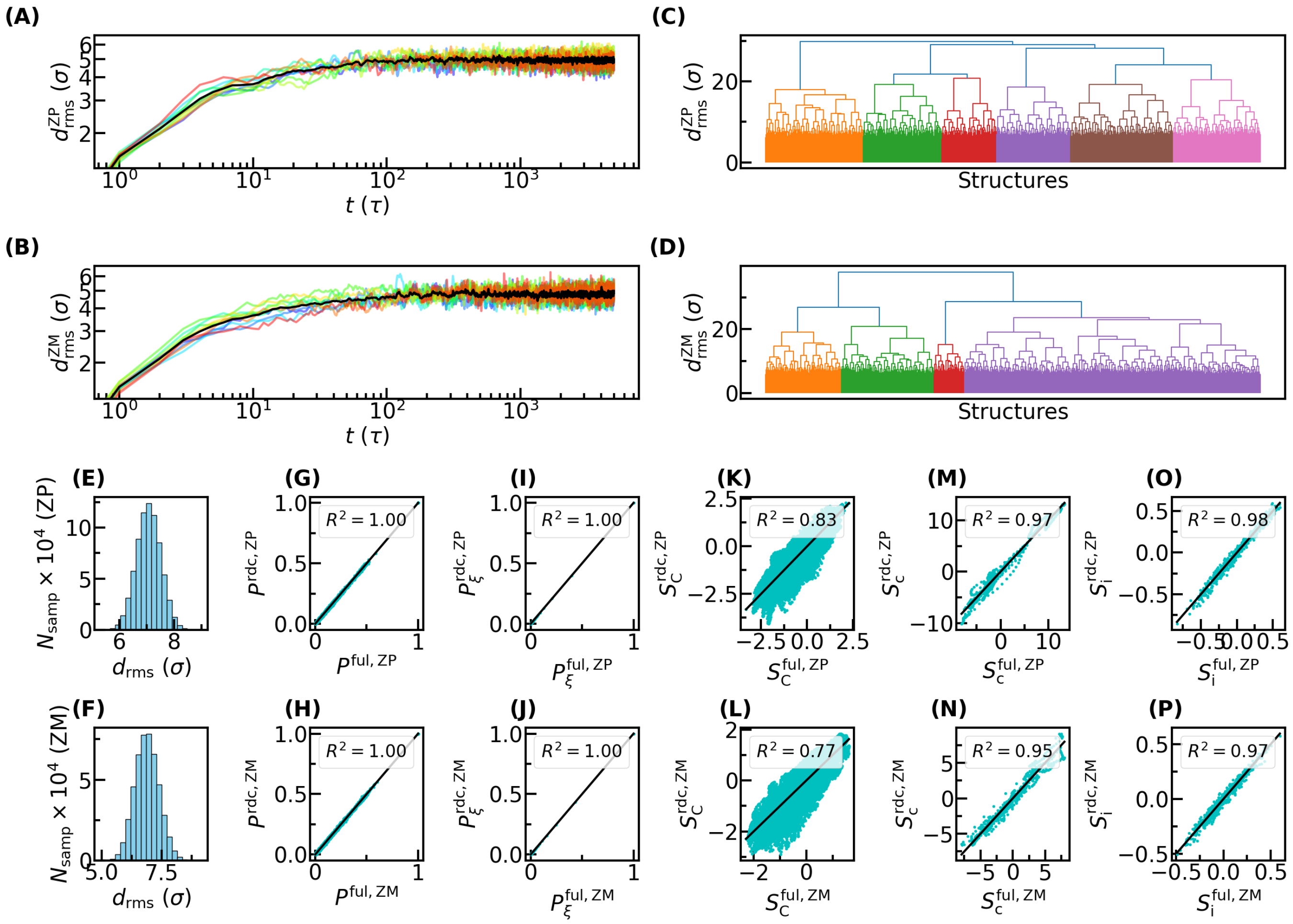


Figure S4: Reduction of initial chromosome structure ensemble for performing landscape-switching simulations. (A) Accessibility range of chromosome structures of ZP during the equilibration diffusion, i.e., how much $d_{\mathrm{rms}}$ can it deviate from its initiate structure after enough time. The colored curves represent 10 typical structure, and the black curve represents the average. (B) Similar to (A), but for ZM. (C) Hierarchical cluster of the raw chromosome structural ensemble of ZP. (D) Similar to (C), but for ZM. (E) Distribution of pair-wise $d_{\mathrm{rms}}$ between the chromosome structures in reduced ensemble of ZP. (F) Similar to (E), but for ZM. (G) Similar to Figure S1B, but for comparison between the contact probability matrices from raw ensemble and reduced ensemble. (H) Similar to (G), but for ZM. (I-P) Similarly, but for comparisons of other indicators

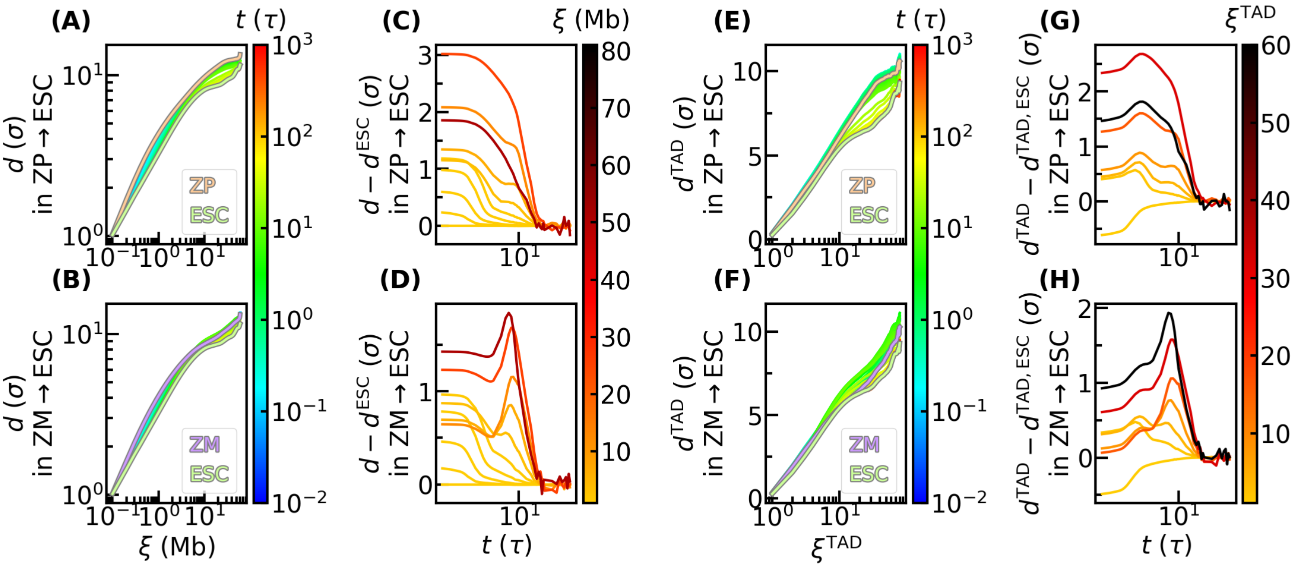


Figure S5: Complementary presentation of Figure 2J-K and Figure 5G-H. (A-B) The x-log-scale presentation of Figure 2J-K. (C-D) Similar to Figure 2J-K, but taking time as x-axis and taking the case of ESC as the zero reference. (E-H) Similar to (A-D), but for Figure 5G-H

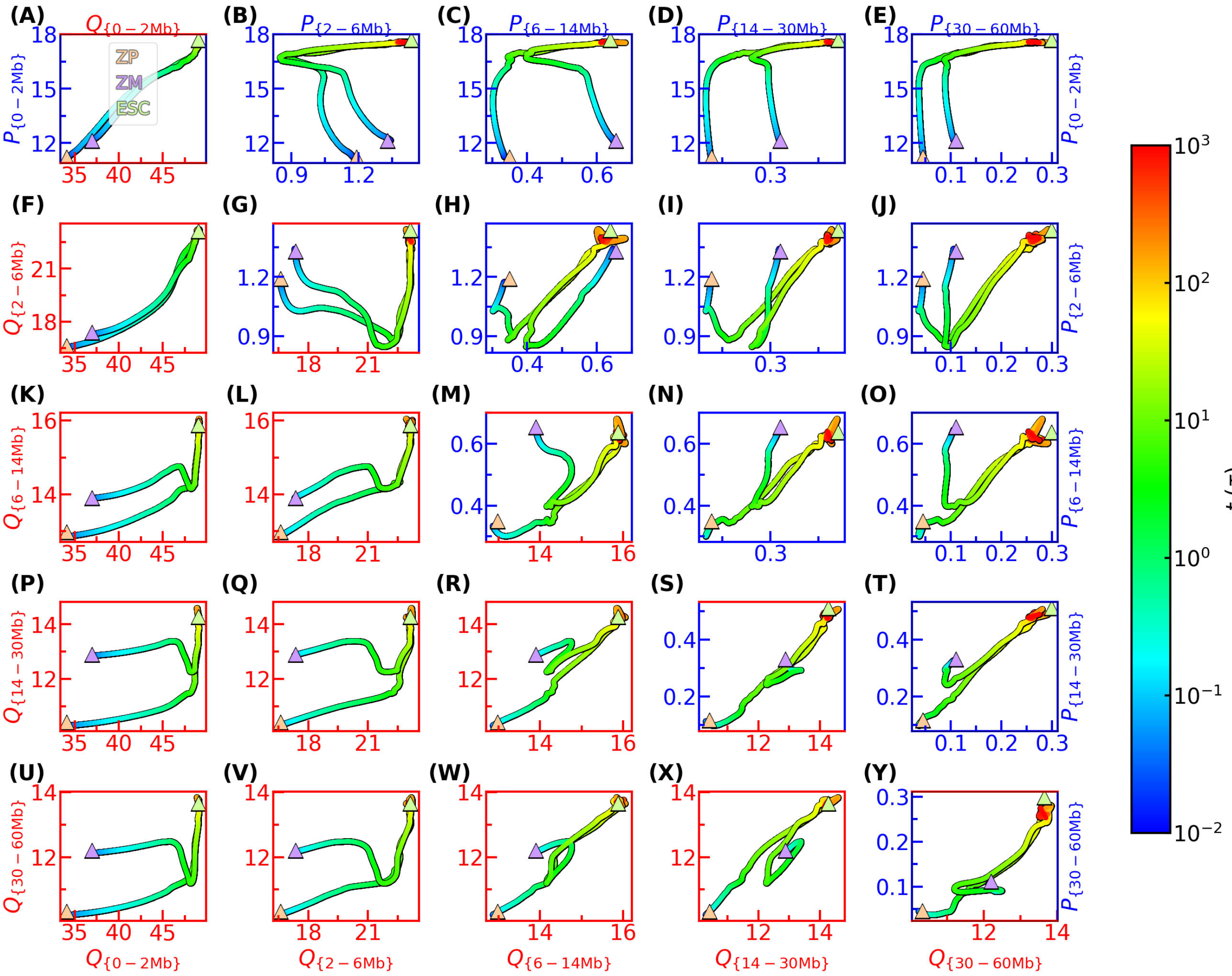

Figure S6: Two dimensional projection of chromosome compactness and folding progress in different length scales. (A) Chromosome compactness and folding progress both in the range of 0–2 Mb. The triangles with gray border represent the value of reference cells. (B-Y) Similar to (A), but for different length scales. The axis colored by blue represents the chromosome compactness in the length scale labeled in this row or column. The axis colored by red represents the chromosome folding progress in the length scale labeled in this row or column.

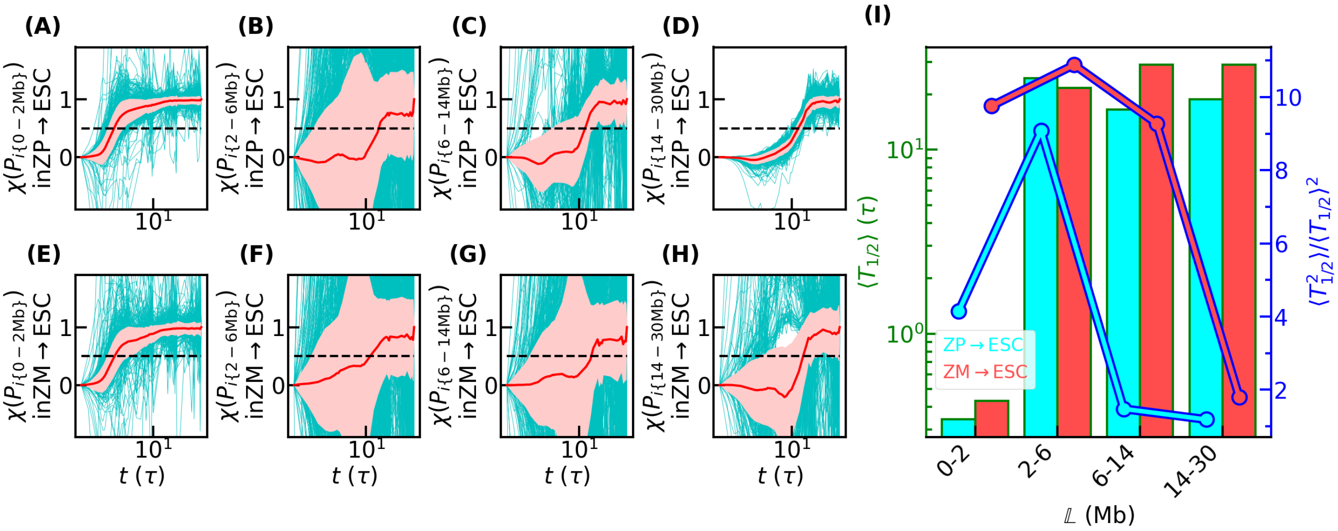


Figure S7: Kinetics, quantified by chromosome compactness $P_{i\mathbb{L}}$, of cell state transition during embryogenesis. (A) Transition pathways of the normalized compactness in the range of 0–2 Mb, for the case of paternal chromosome. The cyan curves are the chromosome compactness formed by each locus with other loci. The red curve is the mean value. The pink shadow indicates the standard deviation. The black dashed line corresponds to the y-value of 1/2. (B-D) Similar to (A), but for 2–6 Mb, 6–14Mb, and 14–30 Mb, respectively. (E-H) Similar to (A-D), but for maternal chromosome. (I) The mean (left y-axis) and specific second-order moment (right y-axis) of the half lives of the transition kinetics quantified by compactness.

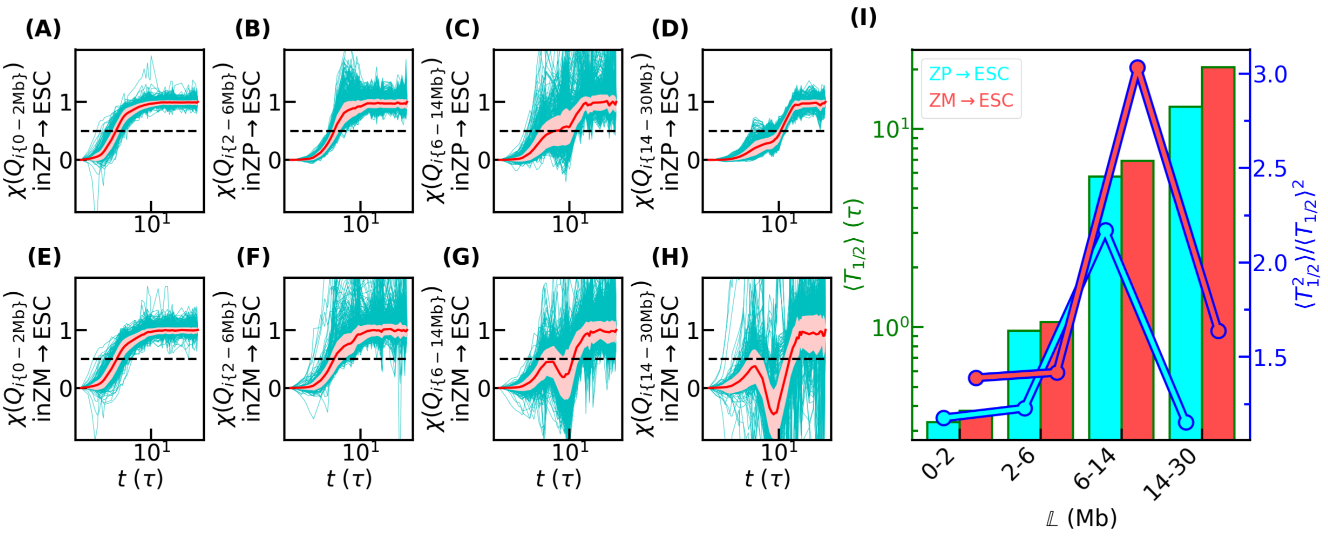

Figure S8: Similar to Figure S7, but for those quantified by folding progress $Q_{i\mathbb{L}}$.

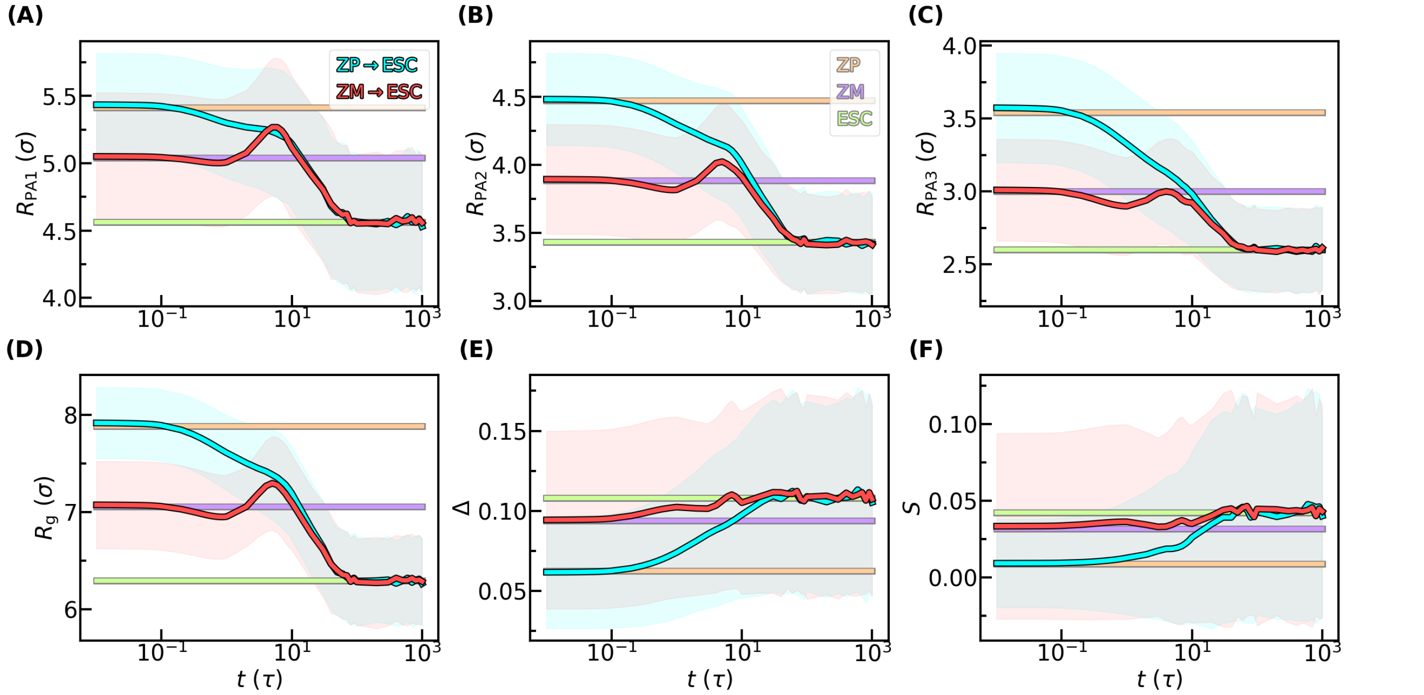


Figure S9: Time evolution of mean lengths of inertial principal axes of (A-C), mean radius of gyration (D), and mean aspherical shape parameters (E-F) of chromosome structures, during embryogenesis. The lines with gray border represent the values of reference cell. The shadows represent the standard deviation.

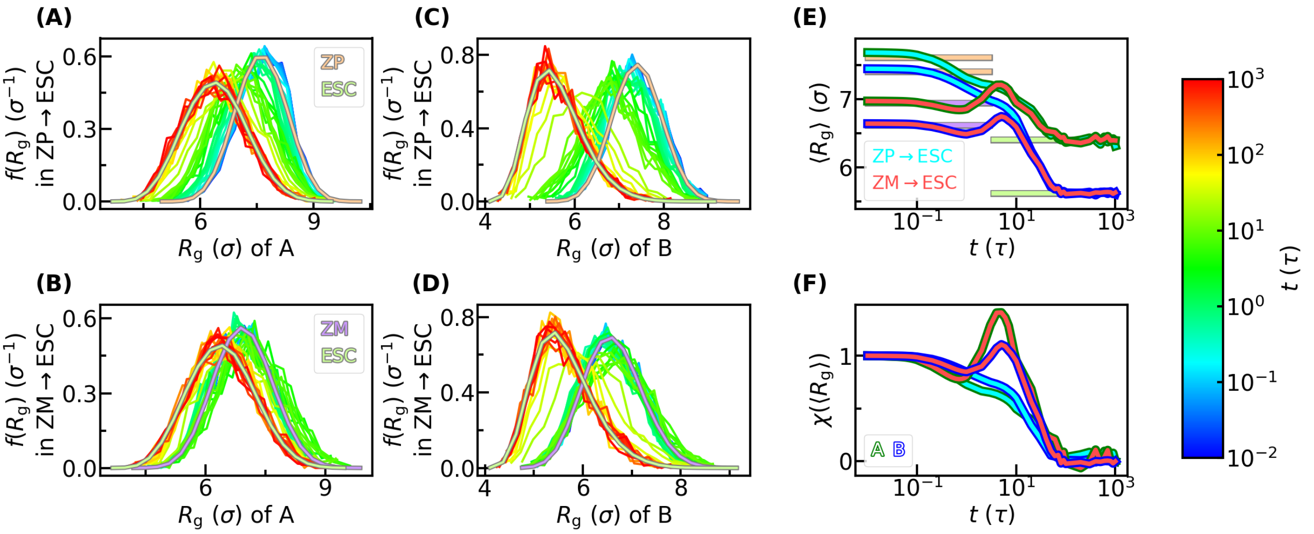


Figure S10: Geometric change of two compartments of chromosomes during embryogenesis. (A) Adaption of distribution of the radius of gyration of compartment A of paternal chromosome. The curves with gray border represent the values of reference cells. (B) Similar to (A), but for maternal chromosome. (C-D) Similar to (A-B), but for compartment B. (E) Time evolution of mean value of (A-D). The lines with gray border represent the values of reference cells. (F) Normalized results of (E) by the $\chi$ function.

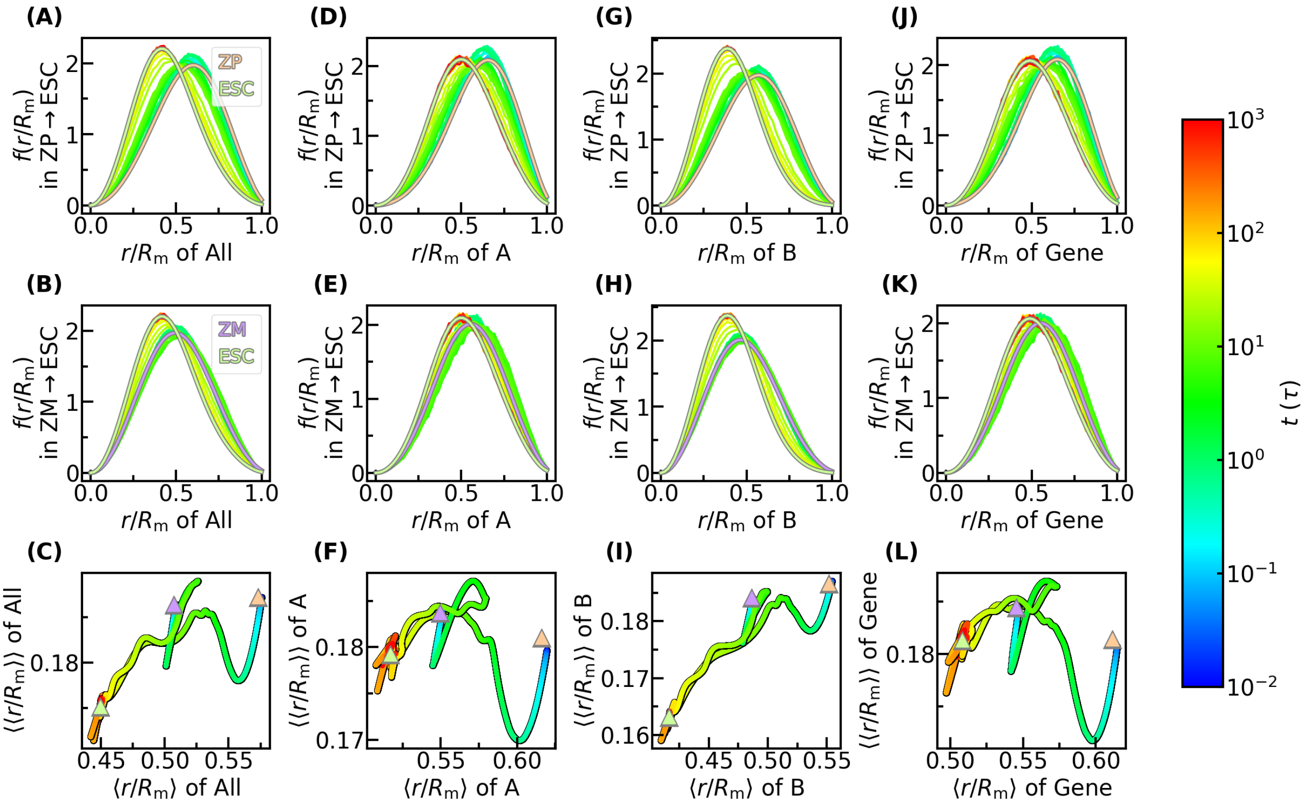

Figure S11: Adaption of chromosomal loci distribution during embryogenesis. (A) Adaption of distribution of absolute radial position of all loci for the case of paternal chromosome. The curves with gray border are the distributions of reference cells. (B) Similar to (A), but for maternal one. (C) Mean and standard deviation of the absolute radial position. The triangles with gray border represent the values of reference cells. (D-F) Similar to (A-C), but for loci in compartment A. (G-I) Similar to (A-C), but for loci in compartment B. (J-L) Similar to (A-C), but for all loci weighted by carried gene number.

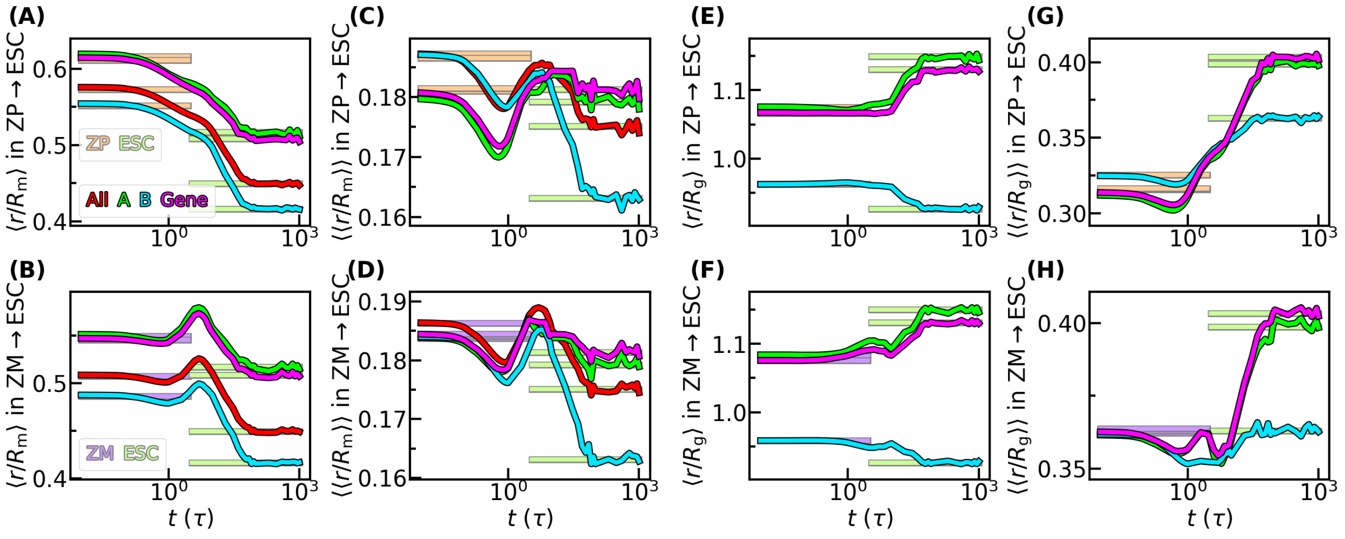


Figure S12: Mean and standard deviation of radial position of loci during embryogenesis in the form of one dimension. (A) Mean absolute radial position of all loci, loci in compartment A, loci in compartment B, and genes, for the case of paternal chromosome. The lines with gray border represent the value of reference cells. (B) Similar to (A), but for maternal one. (C-D) Similar to (A-B), but for standard deviation. (E-H) Similar to (A-D), but for relative radial position.
